# Bivariate genome-wide association study reveals polygenic contributions to covariance between total carotenoid and dry matter contents in yellow-fleshed cassava (*Manihot esculenta*)

**DOI:** 10.1101/2025.03.28.646041

**Authors:** Seren S. Villwock, Elizabeth Y. Parkes, Edwige Gaby Nkouaya Mbanjo, Ismail Y. Rabbi, Jean-Luc Jannink

## Abstract

Cassava breeders aim to increase the provitamin A carotenoid content of storage roots to help combat vitamin A deficiency in sub-Saharan Africa, but a negative genetic correlation between total carotenoid (TC) and dry matter (DM) contents hinders breeding efforts. Genetic linkage between a major-effect variant in the *phytoene synthase 2 (PSY2)* gene and nearby candidate gene(s) has been thought to drive this correlation. Evidence from molecular experiments, however, suggest there may be a metabolic relationship between TC and DM, which we predicted would create genome-wide mediated pleiotropy. Bivariate genome-wide associations were used to test the hypothesis of pleiotropy and examine the genetic architecture of the negative covariance between TC and DM. A population of 378 accessions in the yellow-fleshed cassava breeding program at the International Institute of Tropical Agriculture (IITA) in Ibadan, Nigeria was genotyped with DArTseqLD. TC measured by iCheck™ spectrometer and DM data were available from field trials over ten years across three locations in Nigeria. Mixed linear models controlling for the previously-identified *PSY2* causal variant were used to identify multiple new quantitative trait loci (QTL) jointly associated with both traits. The majority of 17 jointly-associated loci identified at a relaxed significance threshold affected TC and DM in opposite directions, although this pattern did not reach statistical significance in a binomial test. Even after accounting for the effects of these 17 loci as covariates, there was significantly negative polygenic covariance between TC and DM remaining. These findings support the hypothesis that mediated pleiotropy rather than genetic linkage drives the negative genetic correlation between TC and DM in cassava and demonstrate a new application of multivariate GWAS for interrogating the genetic architecture of correlated traits.

**Plain language summary:** Increasing provitamin A in cassava roots has reduced their dry matter content, making vitamin-enriched cassava varieties less desirable. This study used multi-trait models to identify shared genetic factors, most of which had opposing effects on the two traits. The negative relationship was distributed across the genome, suggesting an inherent physiological trade-off. These findings will guide breeders in developing selection strategies for vitamin-enriched cassava and other starchy crops. More broadly, this study demonstrates the use of multi-trait associations to help distinguish whether traits are associated due to separate, nearby genes (genetic linkage) or if the same genes affect multiple traits (pleiotropy).

## Introduction

Vitamin A deficiency is a global public health burden impacting around half of all children in sub-Saharan Africa despite supplementation initiatives (Stevens et al. 2015). Humans require dietary consumption of provitamin A carotenoids to support vision and immune health (Wiseman et al. 2017). Carotenoids are isoprenoid metabolites that are responsible for the yellow, orange, and red pigments in many fruits and vegetables. Certain carotenoids, including β-carotene, α-carotene, and β-cryptoxanthin, are cleaved to form vitamin A in the human body (Perveen et al. 2021). Significant efforts in the last twenty years have focused on biofortifying staple crops with increased carotenoid content to help provide nutritional security for populations at risk for vitamin A deficiencies (Asare-Marfo et al. 2013; Giuliano 2017).

Cassava (*Manihot esculenta* Crantz) has been a particular focus of biofortification efforts as a major food security crop in sub-Saharan Africa, where around 100kg is consumed annually per capita (FAO 2021). Cassava is a woody, tropical plant with starchy storage roots that are a staple food consumed by millions of people globally. Though white-fleshed cassava is generally low in micronutrients, yellow-fleshed varieties accumulate low amounts of provitamin A carotenoids, predominantly β-carotene, which increases both its nutritional value and post-harvest storability (Sánchez et al. 2006; Afolami et al. 2021; Esuma et al. 2021). There are now biofortified varieties of cassava with substantial carotenoid content that have been generated through both conventional selection and genetic engineering (Atser 2014; Beyene et al. 2018; Parkes et al. 2020; Abincha et al. 2024).

In several starchy crops, including cassava and sweetpotato, negative correlations between total carotenoid (TC) content and dry matter (DM) content have been widely reported in breeding populations (Akinwale et al. 2010; Cervantes-Flores et al. 2011; Njoku et al. 2015; Esuma, Kawuki, et al. 2016; Rabbi et al. 2017; Gemenet et al. 2020; Parkes et al. 2020; Esuma et al. 2021). Reductions in DM, which is mainly comprised of starch, negatively impact dry yield, texture, and consumer acceptance of new cassava varieties (Tahirou et al. 2015). It is unclear whether the trade-off between TC and DM is caused by genetic linkage or pleiotropy, with evidence supporting both hypotheses (Ceballos et al. 2013; Rabbi et al. 2017; Beyene et al. 2018; Gutschker et al. 2024; Villwock et al. 2024).

Previous genome-wide association studies (GWAS) in cassava have identified a locus at 24.15 Mbp on chromosome 1 that is associated with both TC and DM, contributing 70% and 37% of the variance for those traits, respectively (Rabbi et al. 2017; Rabbi et al. 2022). This locus contains *phytoene synthase 2 (PSY2)*, the rate-limiting enzyme in carotenoid synthesis, harboring a functionally-validated variant (C_572_A) that increases the catalytic efficiency of PSY2 and promotes carotenoid accumulation (Welsch et al. 2010). There are also two candidate genes for DM at this locus, *sucrose synthase* (SUS) and *ADP-glucose pyrophosphorylase (ADP-glc PPase),* although causal variants in these genes have not been investigated, and the genetic resolution is not high enough to distinguish whether these quantitative trait loci (QTL) are separate from *PSY2*. Similarly, a locus near *PSY2* and *SUS* associated with both TC and DM has also been reported in sweetpotato breeding populations (Gemenet et al. 2020). This evidence strongly supports the hypothesis that an unfavorable genetic linkage at the chromosome 1 locus drives the trade-off between TC and DM. A Latin American cassava population that has overcome the negative correlation through recurrent selection further supports the hypothesis of a breakable genetic linkage (Ceballos et al. 2013). African cassava breeding programs, however, have not been able to replicate these results despite continued selection for both traits.

Other evidence points to a pleiotropic effect of *PSY2* on both TC and DM due to a physiological relationship between the traits. Cassava lines accumulating carotenoids due to expression of *PSY* and *DXS* transgenes exhibited a 50-60% reduction in starch content, along with broad metabolic shifts including increases in sucrose, fatty acid content, and abscisic acid (ABA) content (Beyene et al. 2018). In an African cassava breeding population, yellow-fleshed cassava genotypes exhibited increased biosynthesis of cell wall components, increased sucrose, and decreased starch compared to white-fleshed genotypes (Gutschker et al. 2024). Similar metabolic perturbations have been observed in other crop species engineered to accumulate higher carotenoid content, including potato, maize, and citrus (Diretto et al. 2010; Li et al. 2012; Cao et al. 2015; Decourcelle et al. 2015).

These data suggest the presence of an inherent metabolic interaction between TC and DM, for which several possible biological mechanisms have been proposed, including apocarotenoid signaling and competition for carbon precursors (Olayide et al. 2023; Gutschker et al. 2024; Villwock et al. 2024). If such a metabolic interaction exists, we would expect multiple genomic regions associated with TC to exhibit pleiotropic effects on DM, or *vice versa,* depending on the underlying mechanism. Additionally, we would expect a significant majority of associated genetic variants to affect the traits in opposite directions. Solovieff et al. (2013) define this type of pleiotropy, based on a causal relationship between traits, as “mediated pleiotropy.”

To statistically test the hypothesis of mediated pleiotropy, the identification of more small-effect QTL is needed. Previous GWAS for TC have primarily detected only the *PSY2* locus on chromosome 1 that is responsible for the qualitative difference in carotenoid accumulation between white- and yellow-fleshed cassava, with a few other minor loci were reported on chromosomes 2, 5, 13, and 15 (Esuma, Herselman, et al. 2016; Rabbi et al. 2017; Ikeogu et al. 2019; Rabbi et al. 2022). The yellow-fleshed cassava breeding population at the International Institute of Tropical Agriculture (IITA) in Ibadan, Nigeria presents the opportunity to discover new alleles contributing to quantitative variation in TC and determine if they are jointly associated with DM. Since yellow-fleshed cassava is part of a separate breeding pipeline, there may be different genetic variation within the yellow-fleshed population as opposed to between the white and yellow populations that is overshadowed by the *PSY2* major effect. In addition, the yellow population has been routinely phenotyped using a spectrophotometric device that measures TC more precisely than the indirect estimates of TC in previous GWAS studies, which have used either visual color chart, chromameter measurements, or near-infrared spectroscopy predictions of TC (Jaramillo et al. 2018). Therefore, we genotyped the yellow-fleshed cassava breeding population at IITA and employed a multivariate GWAS approach to test associations between SNPs and the bivariate phenotype of TC and DM.

Multivariate associations have increased statistical power compared to univariate associations when traits are correlated, even if only one phenotype is associated with a given genetic variant (Stephens 2013; Galesloot et al. 2014). This is a counterintuitive but potentially helpful property of multivariate analysis when dealing with an unfavorable correlation between traits because it could facilitate the identification of useful loci that only affect one trait without negatively impacting the other. In some cases, QTL that are not detectable in univariate associations can be identified in multivariate associations due to this increased power (Stephens 2013).

A major limitation of most multivariate GWAS methods is the difficulty in interpreting a significant association with multiple traits, since it is not clear which traits are associated and whether the associations are direct or indirect (Solovieff et al. 2013; Stephens 2013). A common approach is to follow multivariate tests with multiple univariate tests to help interpret which of the traits are associated (Stephens 2013; Taraszka et al. 2022). This approach has limitations because it loses the statistical power benefit of the multivariate test and fails to account for the trait correlations. As a result, the univariate tests may result in false-positive associations with traits that are only indirectly associated with the variant via their correlation with a directly associated trait (Stephens 2013). Our approach overcomes these challenges of traditional multivariate GWAS by explicitly estimating the effect of the SNP on each trait and conducting a bivariate test for joint associations, which facilitates interpretation of how effects on each trait contribute to a multivariate association.

We further expand on a traditional multivariate GWAS. Rather than focusing on single associations, we examine the pattern of associations across the genome to help infer the biological relationship between correlated traits. Specifically, we assess the distribution of jointly-associated SNPs across the genome to test the hypothesis that TC and DM are genetically correlated only because of the *PSY2* major effect locus. If this major locus is the sole or primary cause, we expect that after controlling for the *PSY2* locus and the traits’ overall genetic correlation, other loci would only be associated with a single trait and that their effect directionality would be random. Alternatively, if TC and DM are genetically correlated due to an inherent metabolic or physiological relationship, we expect that most QTL associated with one trait would also be associated with the other trait and would affect the traits in opposite directions. Multivariate models further present a framework for characterizing the genome-wide contributions to the genetic covariance between traits.

To distinguish between mediated pleiotropy versus “biological” or direct pleiotropy, statistical mediation analyses have been used to infer causal relationships between traits (Mackay and Anholt 2024). With mediation analysis, a regression for one trait is fit conditional on an associated trait to test for conditional independence and to partition causal effect into direct and mediated effects (Rockman 2008). Stephens (2013) presented a Bayesian model comparison framework for determining whether multiple correlated traits that are jointly associated with a genetic variant are individually associated directly, indirectly, or unassociated. By using Bayesian model averaging rather than directly fitting each linear model for comparison, the computational time is greatly reduced, but the proposed algorithm did not allow for the correction of population structure and genomic relatedness through covariates. To overcome this challenge, we implemented a frequentist version of the model comparison method described in Stephens (2013) to distinguish between indirect and direct trait associations while properly accounting for population structure.

While these statistical association models may not necessarily reflect biological mechanisms, examining the pattern of associations across the genome provides a framework to evaluate evidence for biological or mediated pleiotropy as contrasted with genetic linkage. Determining whether genetic linkage or pleiotropy is involved will inform breeding strategies for increasing or maintaining both TC and DM, which is critical for the adoption of PVA biofortified staple crops and shed light on potential metabolic interactions between carotenoids and carbohydrates.

## Methods

### Germplasm and genotypic data

A sample of 380 cassava accessions were selected to represent the yellow cassava breeding population at the International Institute of Tropical Agriculture (IITA) in Ibadan, Nigeria, including the HarvestPlus (https://www.harvestplus.org/) and the NextGen Cassava project (https://www.nextgencassava.org/) breeding pipelines. This population originated from crosses between high carotenoid varieties from Latin America and Africa with high-yielding African varieties. Since the early 2000s, recurrent phenotypic selection and marker-assisted selection have been used to improve yield, disease resistance, nutritional quality, and other agronomic traits in the yellow cassava population, which has led to the release of nine yellow varieties since 2011 (HarvestPlus 2024). The population sample used here included recent ancestry from Latin America, with 26 Latin American clones and 34 F1 or F2 hybrids between Latin American and African clones (Fig. S1). 78% of the clones were early-stage selections, while the remainder were more advanced selections. 92% of the clones were classified as “cream” or “yellow” flesh types while the remainder were “white” based on visual color rating. Many accessions of this population have been described and studied previously (Akinwale et al. 2010; Udoh et al. 2017; Ige et al. 2022; Mbanjo et al. 2024; Anokye et al. 2025). Additional accession metadata are available in Table S1.

Leaf tissues from the clones were sampled and sent to Intertek (Parkside, SA, Australia) for DNA extraction. DNA was sequenced, aligned to cassava reference genome v6.1, and 12,981 SNPs were called using the DArTSeqLD platform (Diversity Arrays Technology, Bruce, ACT, Australia). These data were combined with existing genotype data for 50 accessions downloaded from Cassavabase (https://cassavabase.org/). The identities of 9 pairs of duplicate genotype records for the same accession were checked with ‘bcftools gtcheck’ (*bcftools* version 1.18) and highly similar duplicates were removed. Dissimilar duplicates with a high number of mismatching genotypes were both removed, excluding two accessions from further analysis.

To increase mapping resolution, DArTSeqLD SNP data were imputed to a higher density with Beagle5.0 (Browning et al. 2018) using the R package genomicMateSelectR version 0.2.0 (Wolfe 2022) with a reference panel of 21,856 accessions available on Cassavabase (https://cassavabase.org/ftp/marnin_datasets/nextgenImputation2019/) as in Nandudu et al. (2024). A principal component analysis (PCA) showed that the study population clustered within the imputation reference panel, indicating the reference panel appropriately captures the genetic diversity in the study population (Fig. S2). Then, 7,096 accessions available on Cassavabase with both DArTSeqLD genotypes and KASP marker genotypes for the *PSY2* causal allele at S1_24155522 were joined and phased with Beagle5.0 to create a reference panel for imputing the *PSY2* marker (observed directly in 46 accessions). Following imputation, SNPs were filtered with the following parameters: dosage *R*^2^ (DR2) > 0.75 (with the exception of the *PSY2* marker with DR2 = 0.74), Hardy-Weinberg equilibrium p-value (P_HWE) > 1e-20, and minor allele frequency (MAF) > 0.05 for a total of 43,720 biallelic SNPs for 378 unique accessions. Imputed allele dosage expectations were left as unrounded decimals to account for imputation uncertainty (Guan and Stephens 2008; Kutalik et al. 2011).

Population structure was visualized with PCA in R using *prcomp()* on the genetic data matrix and *ggplot()* for plotting. The first five principal components were used to account for population structure in the genome-wide associations based on the elbow of the scree plot, where the proportion of variance explained by PCs began to plateau. An additive genomic relationship matrix calculated using the *rrBLUP* package (Endelman 2011) was used to account for relatedness among the genotypes. Linkage disequilibrium (LD) patterns in the data were evaluated by calculating R^2^, the squared Pearson correlation coefficient, between each pair of SNPs with *cor().*

### Phenotypic data

Phenotypic data were downloaded from Cassavabase (https://cassavabase.org/) for field trials that included at least 3 of the genotyped accessions with carotenoid measurements to ensure sufficient data for estimating trial-level effects. A total of 51 trials included 4 diversity collection trials (“Genetic Gain”; GG), 6 Clonal Evaluation Trials (CET), 17 Preliminary Yield Trials (PYT), 12 Advanced Yield Trials (AYT), 1 Uniform Yield Trial (UYT), and 11 Nationally Coordinated Research Program variety trials (NCRP), representing different stages of selection in the breeding pipeline and levels of replication (Table S2).

Dry matter (DM) content percentage was measured as the ratio of the weight before and after oven drying about 100g of shredded cassava mixed from 8 peeled roots as previously described (Rabbi et al. 2017). Total carotenoids were measured by spectrophotometry using the iCheck™ Carotene device (TCICHK) from BioAnalyt (Teltow, Germany) as previously described (Rodriguez-Amaya and Kimura 2004; Maroya et al. 2012; Udoh et al. 2017; Jaramillo et al. 2018). Briefly, a blended cassava root sample was mixed with an alcohol-based extraction reagent and placed into the iCheck™ spectrophotometer to estimate the concentration of total carotenoids (in *μ*g/g fresh weight (FW)) based on light absorbance. The total carotenoid content in cassava roots is comprised predominantly (∼85%) β-carotene with lower levels of α-carotene and xanthophylls such as violaxanthin and lutein (Rodriguez-Amaya and Kimura 2004; Ikeogu et al. 2017). The iCheck™ method has previously shown a high correlation (r^2^ = 0.98, *p* < 0.001) with HPLC measurement of total carotenoids (Jaramillo et al. 2018). Carotenoid content was also rated by visual color evaluation using a color chart (TCHART) on an ordinal scale from 1 (white) to 7 (deep yellow) (Rabbi et al. 2017). Additional agronomic traits measured for at least 300 accessions were selected for association analyses (Table 1).

**Table 1.** Description, sample size, and broad-sense Heritability (H^2^) of traits analyzed.

| Trait name | Abbreviation | Trait ontology ID | No. of genotypes | No. of observations | H<sup>2</sup> |
| --- | --- | --- | --- | --- | --- |
| Total carotenoids by iCheck™ method | TCICHK | CO_334.0000162 | 365 | 1221 | 0.49 |
| Total carotenoids by color chart (1-9) method | TCHART | CO_334.0000161 | 374 | 1750 | 0.73 |
| Dry matter content percentage | DM | CO_334.0000092 | 370 | 1714 | 0.50 |
| Dry yield | DYLD | CO_334.0000014 | 332 | 1249 | 0.21 |
| Fresh storage root weight per plot | RTWT | CO_334.0000012 | 373 | 1748 | 0.26 |
| Harvest index | HI | CO_334.0000015 | 368 | 1408 | 0.43 |
| Sprouting proportion | SPRTPERC | CO_334.0000008 | 375 | 1697 | 0.24 |
| Fresh root yield | FRYLD | CO_334.0000013 | 334 | 1333 | 0.23 |
| Cassava mosaic disease severity 3-month evaluation | CMD3S | CO_334.0000192 | 376 | 1792 | 0.60 |
| Cassava mosaic disease incidence 3-month evaluation | CMD3I | CO_334.0000196 | 376 | 1756 | 0.49 |

The phenotypic data were analyzed in a two-step approach to account for environmental effects before running the GWAS. Phenotypic values were transformed using the Box-Cox transformation for normality using the R function *boxcox,* using the optimum *λ* value for each trait (Box and Cox 1964). The data were then centered and scaled by dividing by the traits’ standard deviations. Best linear unbiased predictions (BLUPs) of the effect of genotype ID on each trait and variance components were calculated using the following mixed model fit with ASReml-R version 4.1.0.110 (The VSNi Team 2023):

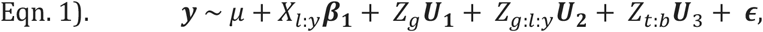

where ***y***is a vector (*n* x 1) of Box-Cox normalized and scaled trait values for *n* observations; *μ* is the overall population mean; *X*_l:*y*_ is a design matrix (*n* x *n*_*ly*_) indicating the year within location for each observation; ***β*_1_** is a vector (*n*_*ly*_ x 1) of fixed effect estimates for year nested within location; *Z*_*g*_ is a design matrix (*n* x *n*_*g*_) indicating the genotype ID for each observation; ***U*_1_** is a vector (*n*_*g*_ x 1) of predicted random effects of genotype (BLUPs) for *n*_*g*_ genotypes, assumed to be normally distributed 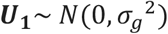 with genetic variance 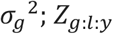 is a design matrix (*n* x *n*_*gly*_) indicating the genotype, location, and year combination for each observation; ***U*_2_** is a vector (*n*_*gly*_ x 1) of predicted random effects of the interaction of genotype with year/location combination, assumed to be normally distributed 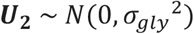 with genotype by environment variance 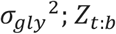 is a design matrix (*n* x *n*_*tb*_) indicating the block number within trial for each observation; ***U*_3_** is a vector (*n*_*tb*_ x 1) of predicted random effects for block within trial, assumed to be normally distributed 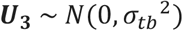 with block variance *σ*_*tb*_^2^; and ***ϵ*** is the vector (*n* x 1) of residual effects, assumed to be normally distributed 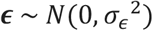 with residual variance 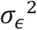. Outliers were identified and removed based on studentized deleted residuals with a Bonferroni corrected α = 0.05 (Kutner et al. 2005). Then, Eqn. 1 was re-fitted without the outliers (4 observations excluded for TCICHK; 3 for TCHART; and 6 for DM). The broad-sense heritability (H^2^) of each trait was calculated as follows: 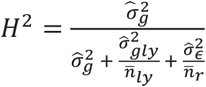, where 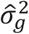 is the estimated genetic variance, 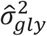 is the estimated genetic by environment variance, 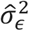 is the estimated residual variance, 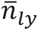 is the harmonic mean number of location/year combinations per accession, and 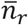 is the harmonic mean number of plots per accession across locations, years, and replications (Holland et al. 2002).

Finally, BLUPs were de-regressed by dividing by their prediction reliability 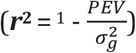 as described in Garrick et al. (2009). To account for the imbalanced number of observations per genotype, the BLUP prediction reliabilities were carried forward to use as inverse variance weights in the second-stage GWAS model as recommended by Garrick et al. (2009). Trait-specific weights were calculated as 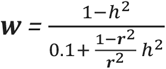, where 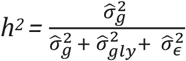.

### Statistical methods for genome-wide associations

Genome-wide associations for the bivariate phenotype were tested using the following mixed linear model fit for each SNP:

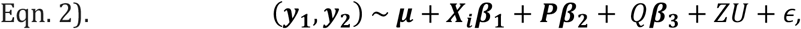

where (***y*_1_, *y*_2_**) is a matrix (*n*_*g*_x 2) of de-regressed TCICHK and DM BLUP values for *n*_*g*_genotypes; ***μ***is a vector (2 × 1) of overall means for each trait; ***X*_*i*_** is a vector (*n*_*g*_x 1) of marker dosages (0, 1, 2) for SNP *i* for *n*_*g*_genotypes; ***β*_1_** is a vector (1 × 2) of estimates for the additive fixed effect of SNP *i* on each trait; ***P*** is a vector (*n*_*g*_x 1) indicating the imputed genotype at the *PSY2* causal SNP, S1_24155522; ***β*_2_** is a vector (1 × 2) of estimates for the additive fixed effect of the *PSY2* SNP on each trait; *Q* is a matrix (*n*_*g*_x 5) of the first 5 PCs for the *n*_*g*_genotypes; ***β*_3_** is a matrix (5 × 2) of fixed effect estimates for the first 5 PCs; *Z* is an identity matrix (*n*_*g*_x *n*_*g*_) identifying the genotype, *U* is a matrix (*n*_*g*_x 2) of the genotype random effect on each trait with matrix normal distribution 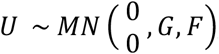, with covariance across rows specified by *G*, the relationship matrix (*n*_*g*_x *n*_*g*_) for the genotypes calculated with genome-wide SNPs excluding the *PSY2* SNP, and covariance across columns specified by *F*, the additive genetic variance-covariance matrix (2 × 2) for the two traits after accounting for the SNP fixed effects; *ϵ* is a matrix (*n*_*g*_x 2) of residual effects for each trait, with matrix normal distribution 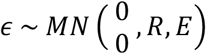 with covariance across rows specified by *R,* a diagonal matrix (n_*g*_x n_*g*_) of inverse BLUP reliability weights ***w***, and covariance across columns specified by *E*, the variance-covariance matrix (2 × 2) of the residual effects for the two traits. The averages of the trait-specific BLUP weights for TCICHK and DM were used as accession-level weights. This equation was fit for each SNP in *ASRemlR* with the following statements:

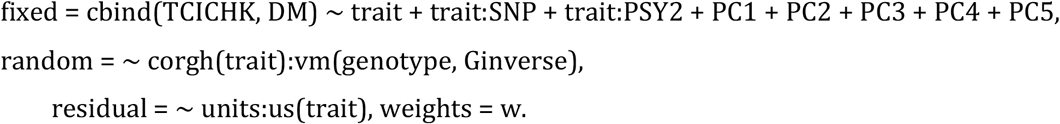

Genome-wide associations for univariate phenotypes were tested using the following mixed linear model fit for each SNP:

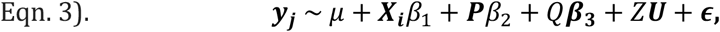

where ***y*_*j*_** is a vector (*n*_*g*_x 1) of de-regressed BLUP values for trait *j* for *n*_*g*_genotypes, *μ* is the overall trait mean; *β*_1_ is the estimated additive fixed effect of SNP *i*; *β*_2_ is the estimated additive fixed effect of the *PSY2* SNP on trait *j* (only included for traits TCICHK, TCHART, and DM); ***β*_3_** is a vector (5 × 1) of fixed effect estimates for the first 5 PCs; ***U***is a vector (*n*_*g*_x 1) of predictions for the genotype random effect, assumed normally distributed 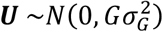 with polygenic genetic variance 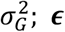 is a vector (*n*_*g*_x 1) of residual effects, assumed normally distributed 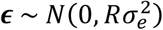 where *R* is a diagonal matrix (*n*_*g*_x *n*_*g*_) of inverse BLUP reliability weights ***w*_*j*_**with residual variance 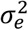 ; and all other variables are as specified above. These models were fit for each SNP and each trait in *ASRemlR* with the following statements:

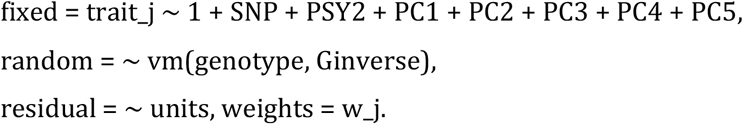

The statistical significance of SNP effects was tested with Wald tests using the R function *aod::wald.test.* SNP effect estimates, and their variance-covariance matrix in the case of the bivariate associations, were extracted from *asreml.predict.* A modified Bonferroni cut-off for α = 0.05 was calculated based on the estimated number of independent tests as described in Gao et al., (2008). The number of eigenvalues explaining 99.5% of the variation in the SNP correlation matrix per chromosome totaled to 3657, which was used as the denominator in the Bonferroni calculation to establish a significance threshold at p < 1.367 × 10^−5^. Manhattan plots were visualized with *ggplot()* and quantile-quantile plots were visualized with the *qqman* package (Turner 2018). A relaxed threshold at p < 0.001 was used to minimize type II error when selecting SNPs for downstream analyses on effect direction. This relaxed threshold enabled the examination of patterns in small-effect loci that could not be identified among the small set of significantly-associated SNPs. While there are likely some false-positive SNPs in this set, we assume they add neutral noise around biological patterns. SNPs with the lowest p-value within a region of high LD (R^2^ > 0.75) were selected to represent independent peaks passing the relaxed threshold. A binomial test was used to determine whether these suggestively-associated SNPs were more likely to affect TC and DM in opposite directions than would be expected by chance.

The proportion variance explained (PVE) by each associated SNP *i* for a given trait was estimated according to the formula 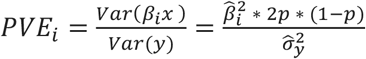, where 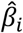 is the estimated additive fixed effect of SNP *i, p* is the frequency of the alternate allele, and 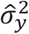 is the total genetic variance, the sum of variance components extracted from the reduced model fit on de-regressed BLUPs without the SNP covariate (Eqn. 2). A measure of proportion covariance explained (PCE) was derived from the PVE formula as follows:

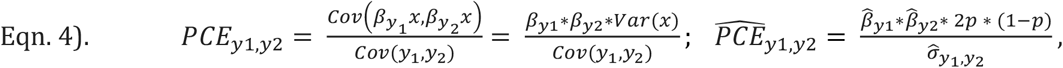

where 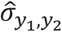 is the estimated total genetic covariance for traits *y*_1_ and *y*_2_, extracted from the reduced model without the SNP. Negative PCE values indicate that the SNP contributes to covariance in the opposite direction of the overall genetic covariance.

### Model comparison

Models specified in Eqn. 2 and 3 were compared with the following mixed linear models using one trait as a covariate for the other:

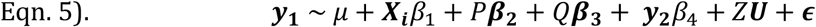

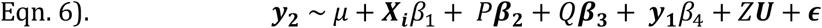

These models were fit for each of the putatively associated SNPs in *ASRemlR* with the following statements:

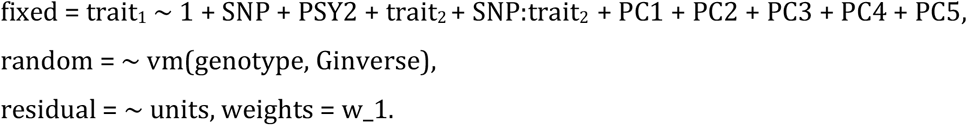

Applying a frequentist version of the framework outlined by Stephens (2013), we used model comparison to determine whether SNPs are directly or indirectly associated with the two traits. Models using trait 2 as a covariate for trait 1 (Eqn. 5) test whether trait 1 is independent of a SNP (*β*_1_ = 0) conditional on trait 2. If a SNP is significantly associated with the bivariate phenotype (Eqn. 2) but is conditionally independent of trait 1 given trait 2 as a covariate, it suggests that the bivariate association is mainly driven by the overall trait correlation, representing a direct relationship with trait 2 and an indirect relationship trait 1 (Fig. 1). Conversely, a SNP that is still associated with trait 1 after accounting for trait 2 as a covariate suggests that the SNP has a direct relationship with both traits. The reasoning is the same using trait 1 as a covariate for trait 2 (Eqn. 6). These possible pleiotropic relationships are summarized in Fig. 1.

**Figure 1.**
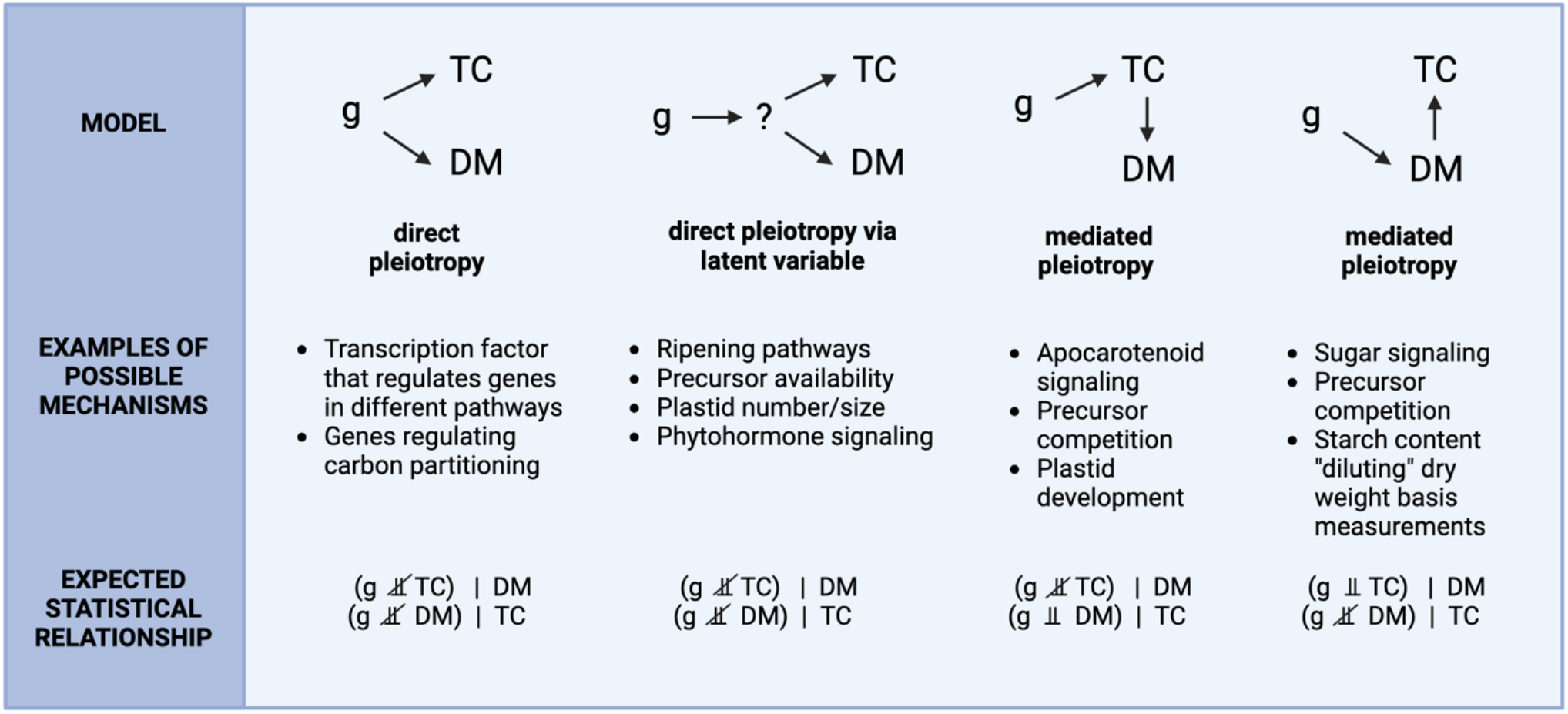
Illustration of direct and mediated pleiotropic relationships and their expected statistical associations, where g ⫫ A | B denotes conditional independence of a genetic variant *g* with trait A given trait B. Hypothesized biological mechanisms are described in detail in Villwock et al. (2024). Created with Biorender.com.

### Estimates of polygenic contribution to negative genetic covariance between TCICHK and DM

The relative contributions of the identified putative QTL to the genetic covariance between TCICHK and DM were assessed by comparing the variance components before and after accounting for the peak SNPs as covariates. A bivariate model as described in Eqn. 2 was fit with all *z* peak SNPs identified at the relaxed significance threshold added as covariates:

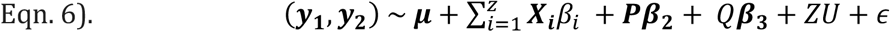

With the genotype random effect 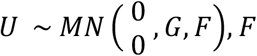 represents the additive genetic variance-covariance matrix for the two traits remaining after accounting for the fixed effects of the identified QTL. If the identified QTL completely accounted for the negative genetic covariance between traits, then the residual genetic covariance in *F* would be expected to approach zero. To formally test whether significant residual genetic covariance remained after accounting for the associated SNPs, a likelihood ratio test implemented with *lrt.asreml()* compared models with the residual genetic covariance between traits set to zero or unconstrained.

### Candidate gene analysis

*M. esculenta* reference genome v6.1 gene annotation data were downloaded from the JGI data portal (https://data.jgi.doe.gov/). A list of *a priori* candidate genes involved in the carotenoid synthesis (MetaCyc pathway ontology ID PWY-6475), apocarotenoid synthesis (PWY-6806), starch synthesis (PWY-622), and starch degradation (PWY-6724) pathways annotated in *Manihot esculenta v6.1* on Phytozome was assembled (Table S3) (Goodstein et al. 2012; Bredeson et al. 2016; Caspi et al. 2020). *A priori* candidate genes were screened for LD and physical distance to significantly associated SNPs. Then, all annotated genes within 100kb of the peak SNPs were extracted from the genome annotation file. Genes with missing ontology annotations were supplemented with annotations from the UniProt Knowledgebase where possible (Bateman et al. 2023).

## Results

### Trait distributions and relationships

The phenotypic distributions of TCICHK and DM observations of genotyped accessions before transformation and statistical analysis are shown in Fig. S3. TCICHK ranged from 0 to 30.8 μg/g FW and DM ranged from 9.1 to 60.0%. TCICHK and DM had moderately high broad-sense heritability of 0.49 and 0.50, respectively. The two traits were negatively genetically correlated with r = -0.34 and p = 1.84 × 10^−11^ (Fig. 2) and phenotypically correlated with r = -0.15 and p = 4.48 × 10^− 22^. TCICHK was also negatively genetically correlated with several other agronomic traits: dry yield (DYLD; r = -0.17, p = 1.66 × 10^−3^), root weight (r = -0.18, p = 5.98 × 10^−4^), fresh root yield (FRYLD; r = -0.15, p = 8.43 × 10^−3^), and sprouting percentage (SPRTPERC; r = -0.25, p = 1.26 × 10^−6^) (Fig. 3). The broad-sense heritabilities of all traits are listed in Table 1.

**Figure 2.**
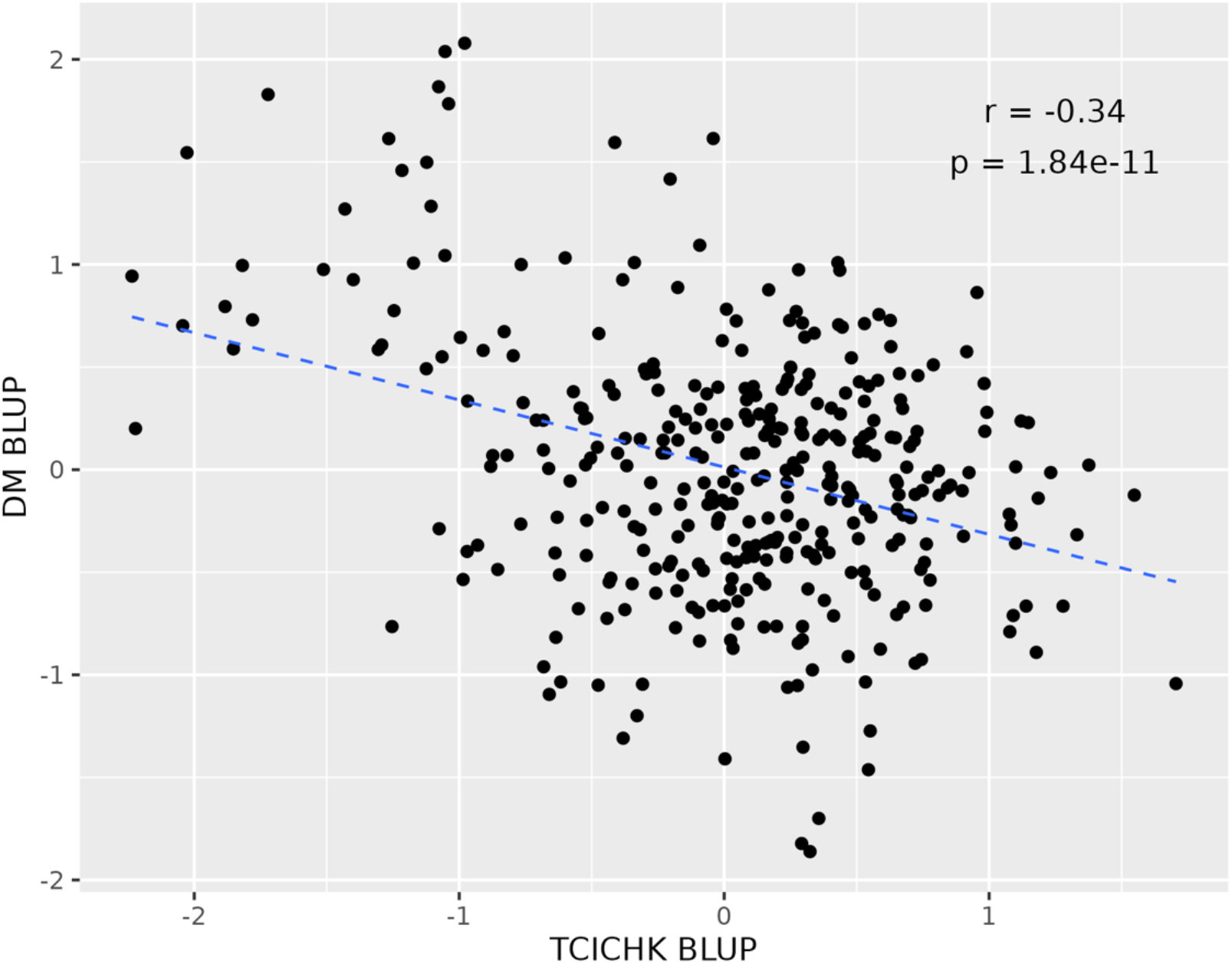
Negative genetic relationship between total carotenoid content as measured by iCheck™ Carotene (TCICHK) and dry matter content (DM) BLUPs among the 378 genotyped accessions after normalization of phenotypic values with Box-Cox transformation. Pearson correlation coefficient (r) and p-value are displayed at the top right.

**Figure 3.**
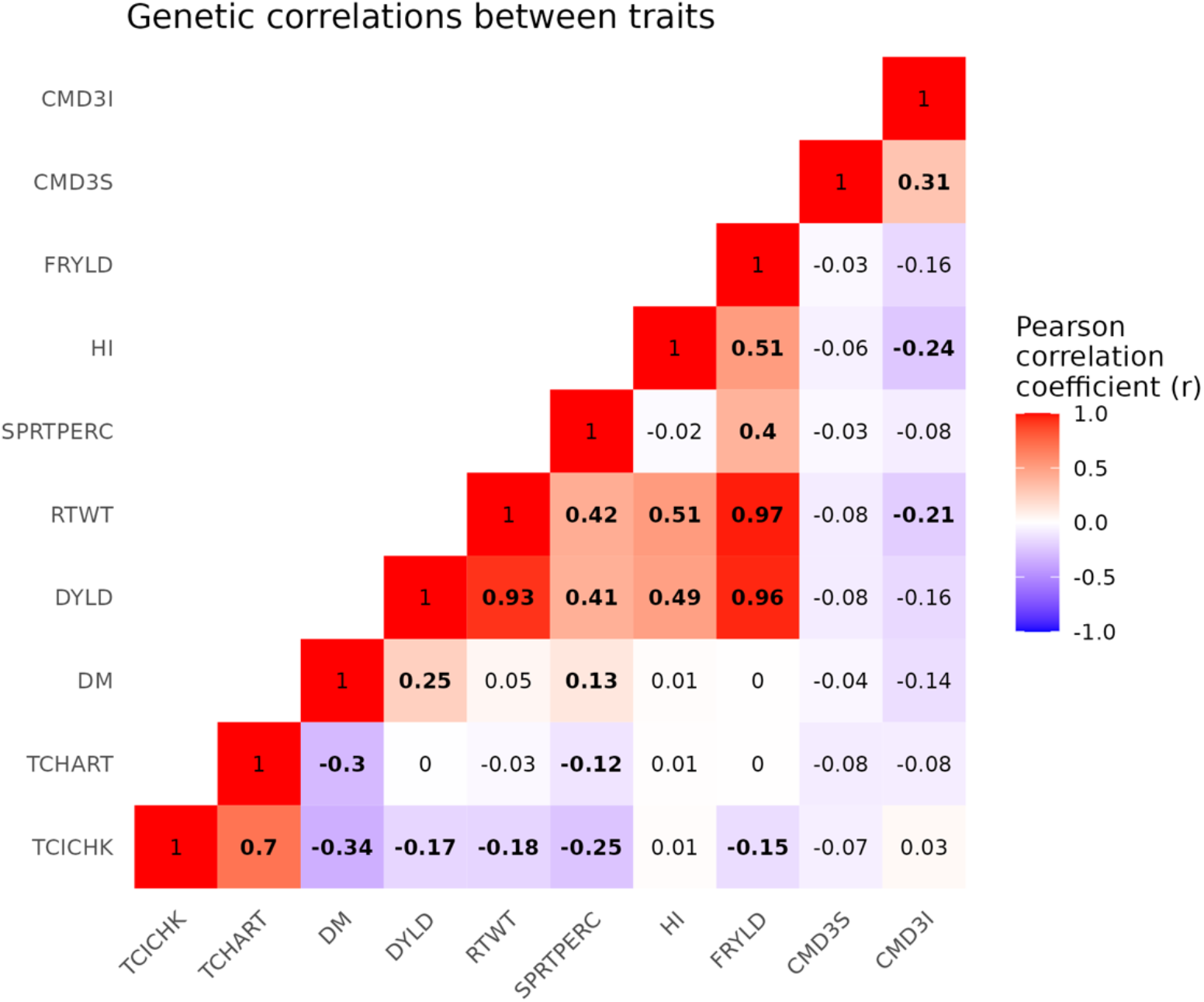
Pairwise genetic correlations between all analyzed traits. Each cell is labeled with the Pearson correlation coefficient (r) between the traits listed on the x and y axis. Significant correlations (p < 0.05) are bolded.

### Bivariate genome-wide associations for TCICHK and DM

The Manhattan plot from the bivariate associations revealed 17 peaks that passed the relaxed threshold (p < 0.001), including two QTL peaks that passed the modified Bonferroni significance threshold for joint association with both TCICHK and DM after controlling for the effect of *PSY2* (Fig. 4). The significant joint associations were located on chromosomes 1 and 10. The top SNP on chromosome 1 is located at 23.484417 Mbp, which is 0.67 Mbp upstream from *PSY2* and was in moderate LD (R^2^ = 0.21) with the *PSY2* causal variant at 24.155522 Mbp. Fig. 5 illustrates the results from bivariate and univariate associations in this QTL region in higher resolution. The top jointly-associated SNP on chromosome 1 in the bivariate associations was significantly associated with TCICHK but not with DM in univariate analyses. The proportion of genetic variance explained by the two QTL were relatively small (PVE_TCICHK_ = 3.4% and PVE_DM_ = 1.1% for the chromosome 1 peak; PVE_TCICHK_ = 2.9% and PVE_DM_ = 0.7% for the chromosome 10 peak). The *PSY2* SNP explained about 2.4% of the genetic variation in TCICHK and 5.0% of the genetic variation in DM. The chromosome 1 and 10 QTL explained about 3% and 2% of the genetic covariance between traits, respectively, while the *PSY2* SNP explained 5.6%. Summary statistics for all associated SNPs are listed in Table 2.

**Table 2.** Summary statistics of SNPs that are jointly-associated with TCICHK and DM. The SNP with the lowest p-value was selected from each group of associated SNPs. P-values are annotated with two asterisks if they pass the modified Bonferroni significance threshold and a single asterisk if they pass the relaxed threshold. SNP effect estimates (additive effect of the ALT allele) and standard errors (SE) are with respect to the transformed, de-regressed BLUP values. PVE and PCE (proportion variance explained and proportion covariance explained, respectively) are with respect to total genetic variance/covariance. Negative PCE values indicate the SNP weakens the overall negative genetic covariance between traits by contributing to covariance the opposite direction.

| Peak SNP<br>(chr#_position) | Bivariate<br>association<br>p-value | TC<br>effect | TC<br>effect<br>SE | DM<br>effect | DM<br>effect<br>SE | Effect<br>direction<br>(TC/DM) | REF<br>allele | ALT<br>allele | Allele<br>freq.<br>(ALT) | TC PVE | DM PVE | TC/DM<br>PCE |
| --- | --- | --- | --- | --- | --- | --- | --- | --- | --- | --- | --- | --- |
| S1_23484417 | 8.81e-06 ** | 0.424 | 0.093 | -0.217 | 0.101 | +/- | C | G | 0.92 | 0.061 | 0.018 | 0.031 |
| S10_2963170 | 1.17e-05 ** | -0.316 | 0.069 | 0.145 | 0.076 | -/+ | A | C | 0.13 | 0.051 | 0.012 | 0.023 |
| S3_3292443 | 4.76e-05 * | -0.022 | 0.072 | -0.310 | 0.071 | -/- | A | T | 0.16 | <0.001 | 0.067 | -0.004 |
| S6_3256937 | 6.38e-05 * | 0.370 | 0.084 | -0.056 | 0.088 | +/- | G | C | 0.06 | 0.035 | 0.001 | 0.005 |
| S13_25853831 | 2.00e-04 * | -0.283 | 0.069 | 0.067 | 0.071 | -/+ | C | T | 0.19 | 0.056 | 0.004 | 0.013 |
| S17_20165942 | 2.66e-04 * | 0.249 | 0.076 | 0.143 | 0.080 | +/+ | C | A | 0.12 | 0.030 | 0.011 | -0.017 |
| S13_23763339 | 2.82e-04 * | -0.217 | 0.058 | 0.116 | 0.060 | -/+ | A | G | 0.29 | 0.044 | 0.014 | 0.023 |
| S5_22756786 | 2.84e-04 * | 0.105 | 0.046 | -0.169 | 0.047 | +/- | A | T | 0.76 | 0.009 | 0.027 | 0.015 |
| S3_294409 | 3.12e-04 * | -0.254 | 0.083 | 0.266 | 0.091 | -/+ | T | A | 0.08 | 0.021 | 0.025 | 0.021 |
| S2_2073445 | 3.43e-04 * | 0.249 | 0.064 | 0.027 | 0.071 | +/+ | C | A | 0.82 | 0.043 | 0.001 | -0.005 |
| S6_23101017 | 3.79e-04 * | -0.368 | 0.094 | 0.004 | 0.102 | -/+ | T | G | 0.06 | 0.035 | <0.001 | <0.001 |
| S17_17210448 | 5.47e-04 * | -0.411 | 0.106 | 0.073 | 0.108 | -/+ | C | A | 0.06 | 0.047 | 0.002 | 0.008 |
| S2_8411339 | 5.65e-04 * | -0.065 | 0.062 | -0.224 | 0.064 | -/- | A | G | 0.21 | 0.003 | 0.043 | -0.011 |
| S10_20300591 | 5.80e-04 * | 0.164 | 0.050 | -0.127 | 0.051 | +/- | T | A | 0.85 | 0.016 | 0.010 | 0.012 |
| S12_5428221 | 6.55e-04 * | -0.013 | 0.053 | 0.205 | 0.054 | -/+ | C | A | 0.38 | <0.001 | 0.051 | 0.003 |
| S18_6776129 | 7.33e-04 * | 0.131 | 0.073 | 0.219 | 0.074 | +/+ | A | C | 0.15 | 0.010 | 0.030 | -0.016 |
| S5_1950967 | 8.67e-04 * | -0.248 | 0.090 | -0.199 | 0.102 | -/- | T | G | 0.08 | 0.020 | 0.015 | -0.016 |

**Figure 4.**
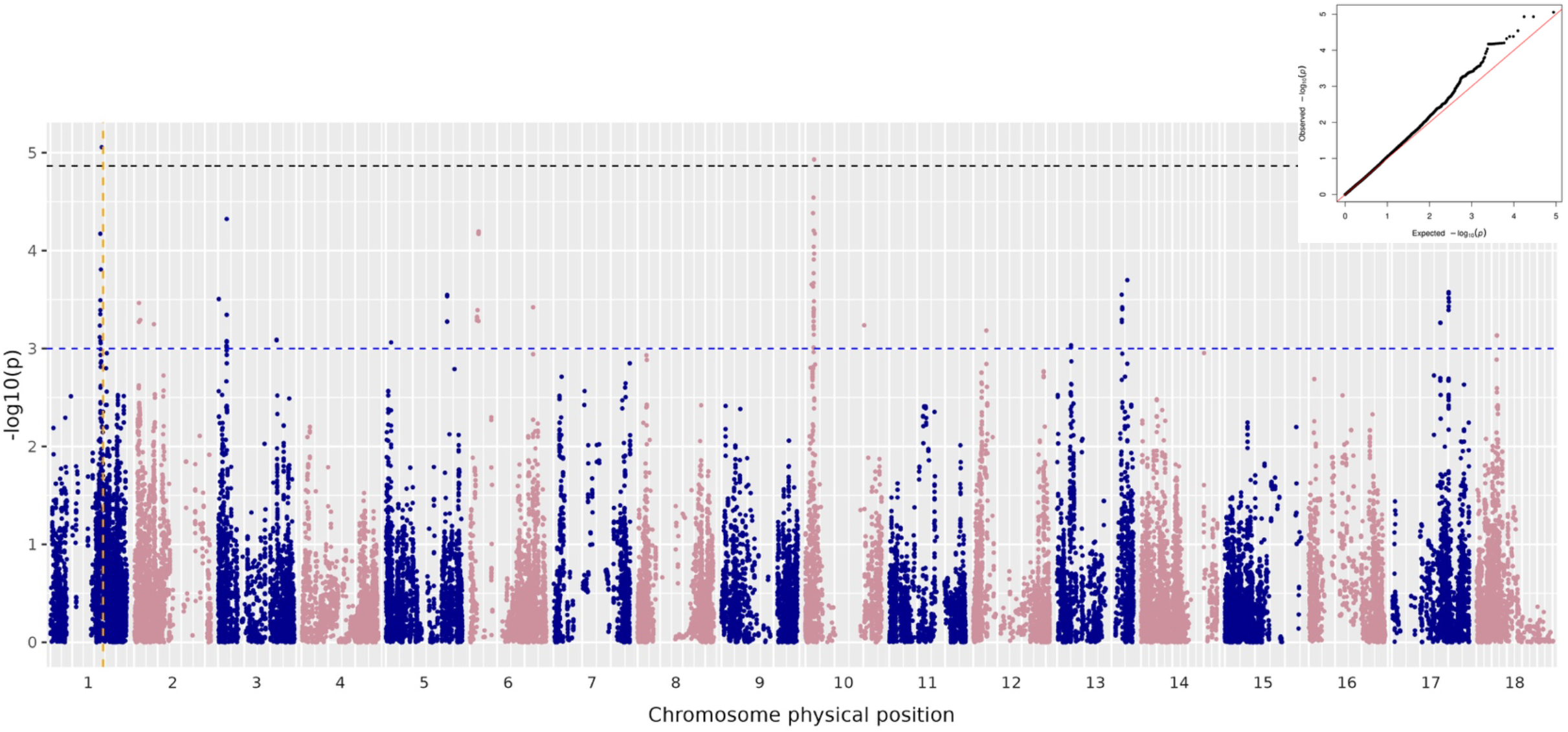
Manhattan plot showing p-values from bivariate Wald tests of SNP effects on TCICHK and DM, accounting for genomic relatedness and the *PSY2* causal SNP. The vertical orange dashed line indicates the position of the *PSY2* causal SNP. The black dashed line indicates the modified Bonferroni significance cut-off at α = 0.05. The blue dashed line indicates a relaxed threshold of p < 0.001 used for downstream analyses. At top right, the Q-Q plot of expected versus observed p-values shows acceptable control of log_10_ p-value inflation.

**Figure 5.**
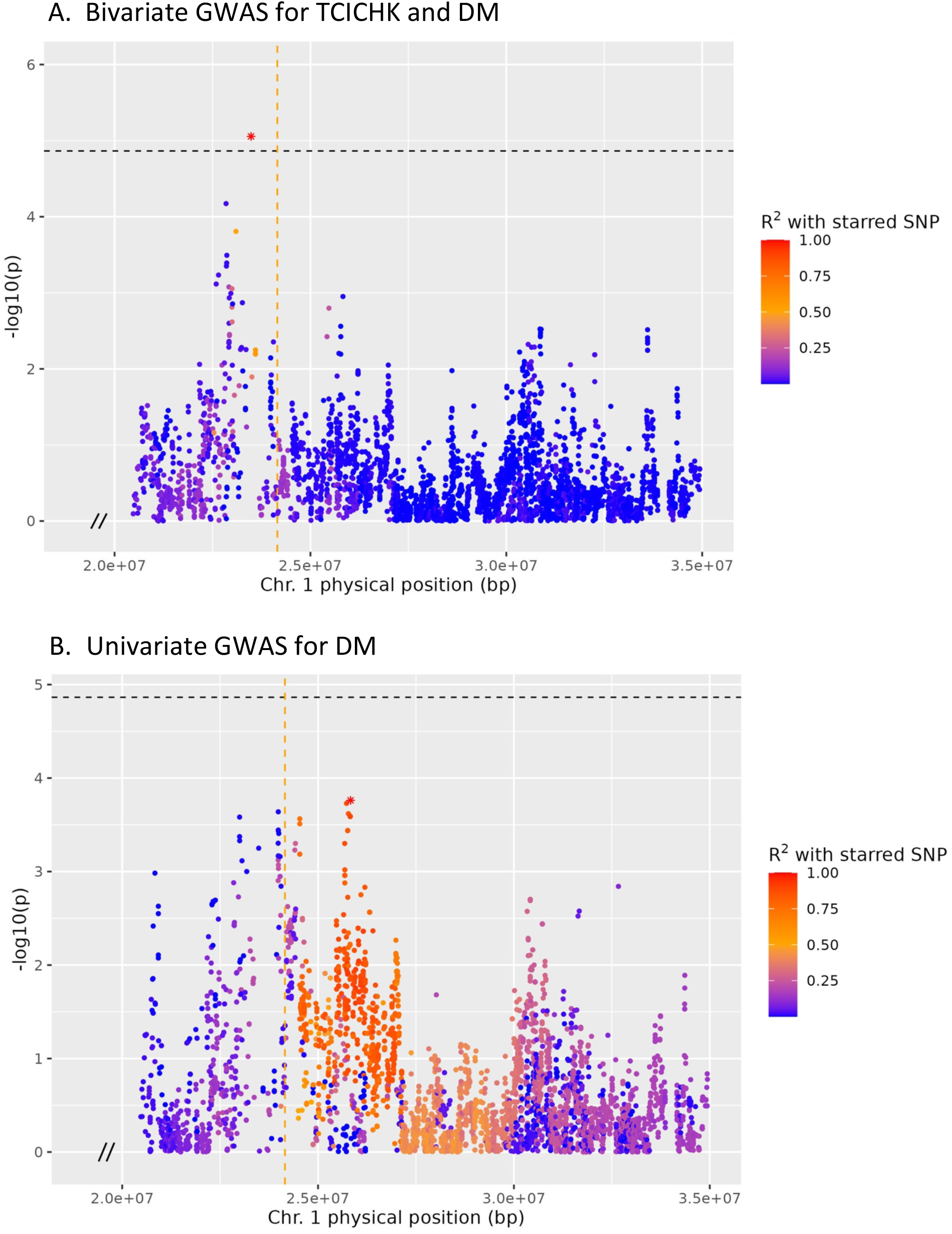
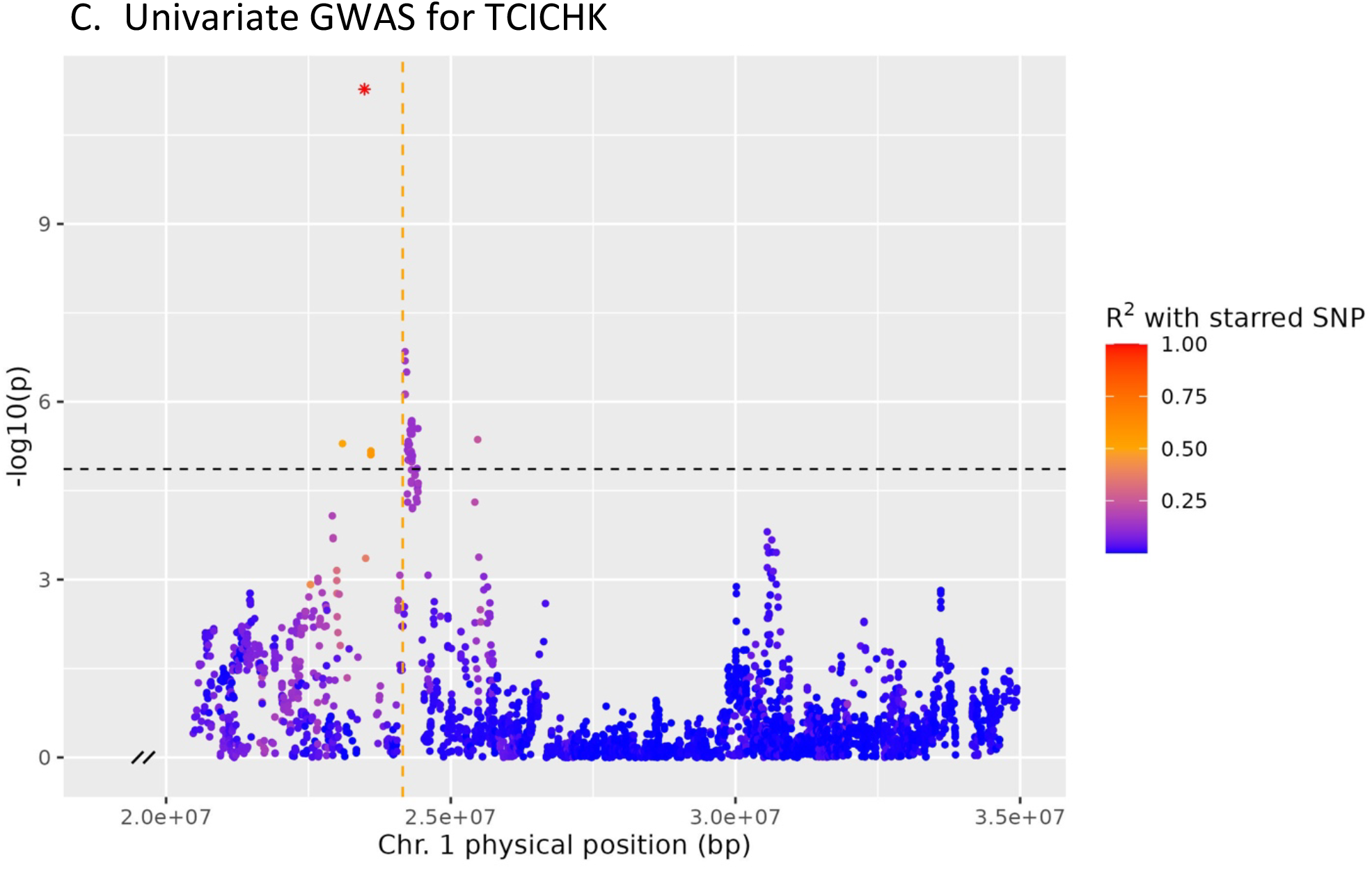
Manhattan plots for A) bivariate TCICHK and DM, B) univariate DM, and C) univariate TCICHK associations on chromosome 1 near the *PSY2* QTL region from 20–35 Mbp. The vertical orange line indicates the position of the *PSY2* causal SNP. Points are colored by their LD (pairwise R^2^) with the top SNP, marked with an asterisk, which differs across association models. The black dashed line indicates the modified Bonferroni significance cut-off at α = 0.05.

Of the 17 associated loci identified at the relaxed significance threshold, 11 peak SNPs had opposite effects on TC and DM, while 6 SNPs had effects on TC and DM in the same direction. A binomial test with these counts resulted in a p-value of 0.166, which was insufficient to reject the null hypothesis that associated SNPs are equally likely to affect TC and DM in same or opposite directions. A likelihood ratio test indicated there was significant negative genetic covariance between TCICHK and DM remaining after accounting for all 17 of these SNPs and *PSY2* as covariates in a linear model (LR-statistic = 9.976, p = 0.0025). Across all SNP association models fit in the bivariate GWAS, the residual genetic covariance was consistently negative, ranging from -0.34 to - 0.19. Together, the peak SNPs accounted for 8.6% and the *PSY2* SNP accounted for 5.6% of the negative covariance between traits.

Since the phenotypes were normalized with a Box-Cox transformation before analysis, bivariate association models (Eqn. 2) were re-fit with untransformed, de-regressed BLUPs for the 17 peak SNPs to translate their effect size estimates to the traits’ original units. For both TCICHK and DM, effect estimates from untransformed data were linearly related (r^2^ > 0.99) to effect estimates from Box-Cox-transformed data with slopes of 5.8 and 6.7, respectively (Fig. S4). SNP effect estimates from the untransformed BLUPs ranged from -2.42 to 2.56 *μ*g/g FW for TCICHK and -1.91 to 1.72% for DM.

### Model comparison

The bivariate association model was compared with univariate models with one trait as the response variable and the other as a covariate to determine whether the associations were direct or indirect. For each of the top SNPs from 17 peaks that passed the relaxed significance threshold, the bivariate model had the lowest AIC compared to the univariate models (Table 3). For 13 of the 17 peak SNPs, the SNP effect retained its significance at the p < 0.001 threshold in a univariate model for one trait with the other as a covariate. For nine of these SNPs, the TCICHK univariate model retained its significance, indicating support for a direct association with TC and indirect association with DM. The remaining four SNPs retained their significance in the DM univariate model, suggesting a direct association with DM and an indirect association with TC. In no cases did the SNP retain significance in both univariate models with the other trait as a covariate.

**Table 3.** Comparison of bivariate and reciprocal univariate association models for the 17 peak SNPs identified as jointly associated at the relaxed threshold (p < 0.001). “TCICHK model” refers to an association between a SNP and TCICHK with DM as a covariate (Eqn. 4), while the “DM model” is the reverse (Eqn. 5). P-values are from univariate or bivariate Wald tests of SNP effects. P-values are annotated with two asterisks if they pass the modified Bonferroni significance threshold and a single asterisk if they pass the relaxed threshold.

| Peak SNP<br>(chr#_position) | Bivariate<br>model AIC | TCICHK<br>model AIC | DM model<br>AIC | Bivariate<br>model p-value | TCICHK<br>model p-value | DM model<br>p-value | Supported model |
| --- | --- | --- | --- | --- | --- | --- | --- |
| 1_23484417 | -47.28 | 3.07 | 0.56 | 8.81e-06 ** | 3.85e-09 ** | 6.25E-03 | Direct TC, indirect DM |
| 10_2963170 | -46.83 | 4.37 | 2.79 | 1.17e-05 ** | 3.86e-05 * | 2.36E-01 | Direct TC, indirect DM |
| 3_3292443 | -44.93 | 22.62 | -12.52 | 4.76e-05 * | 7.81E-02 | 8.81e-06 ** | Direct DM, indirect TC |
| 6_3256937 | -45.12 | 2.62 | 3.47 | 6.38e-05 * | 2.12e-06 ** | 9.44E-01 | Direct TC, indirect DM |
| 13_25853831 | -42.37 | -0.53 | 3.48 | 2.00e-04 * | 1.24e-06 ** | 7.54E-01 | Direct TC, indirect DM |
| 17_20165942 | -42.81 | 5.79 | -0.05 | 2.66e-04 * | 1.19E-03 | 3.76E-03 | NA |
| 13_23763339 | -41.57 | 10.15 | 6.37 | 2.82e-04 * | 1.01e-04 * | 1.56E-01 | Direct TC, indirect DM |
| 5_22756786 | -39.40 | 12.37 | -0.16 | 2.84e-04 * | 6.98E-02 | 3.25E-03 | NA |
| 3_294409 | -42.28 | 13.24 | -4.89 | 3.12e-04 * | 2.69E-03 | 1.66E-02 | NA |
| 2_2073445 | -41.11 | 2.00 | 6.21 | 3.43e-04 * | 1.11e-04 * | 2.80E-01 | Direct TC, indirect DM |
| 6_23101017 | -42.63 | 5.03 | 5.28 | 3.79e-04 * | 3.13e-04 * | 3.41E-01 | Direct TC, indirect DM |
| 17_17210448 | -42.49 | 1.55 | 4.88 | 5.47e-04 * | 3.13e-05 * | 9.20E-01 | Direct TC, indirect DM |
| 2_8411339 | -41.16 | 21.43 | -6.49 | 5.65e-04 * | 1.90E-02 | 7.69e-04 * | Direct DM, indirect TC |
| 10_20300591 | -38.65 | 12.62 | 4.68 | 5.80e-04 * | 4.35E-03 | 1.46E-01 | NA |
| 12_5428221 | -39.56 | 23.14 | -11.18 | 6.55e-04 * | 6.57E-01 | 5.23e-05 * | Direct DM, indirect TC |
| 18_6776129 | -41.06 | 18.37 | -4.00 | 7.33e-04 * | 4.75E-02 | 9.09e-05 * | Direct DM, indirect TC |
| 5_1950967 | -39.50 | 13.10 | -1.26 | 8.67e-04 * | 5.27e-06 ** | 1.24E-01 | Direct TC, indirect DM |

### Univariate associations for all traits

Several peaks passing the modified Bonferroni significance threshold were identified in univariate analyses: TCICHK (7), DM (2), TCHART (4) (Fig. 6). TCICHK and TCHART QTL overlapped on chromosomes 1 and 13. There were no overlapping univariate QTL between TCICHK or TCHART and DM. The peak SNP positions and summary statistics for univariate QTL are reported in Table S4. Among other agronomic traits analyzed (DYLD, FRYLD, RTWT, HI, SPRTPERC, CMD3I, and CMD3S), 2 QTL for SPRTPERC and 1 for CMD3I were identified at the Bonferroni significance level (Fig. S5).

**Figure 6.**
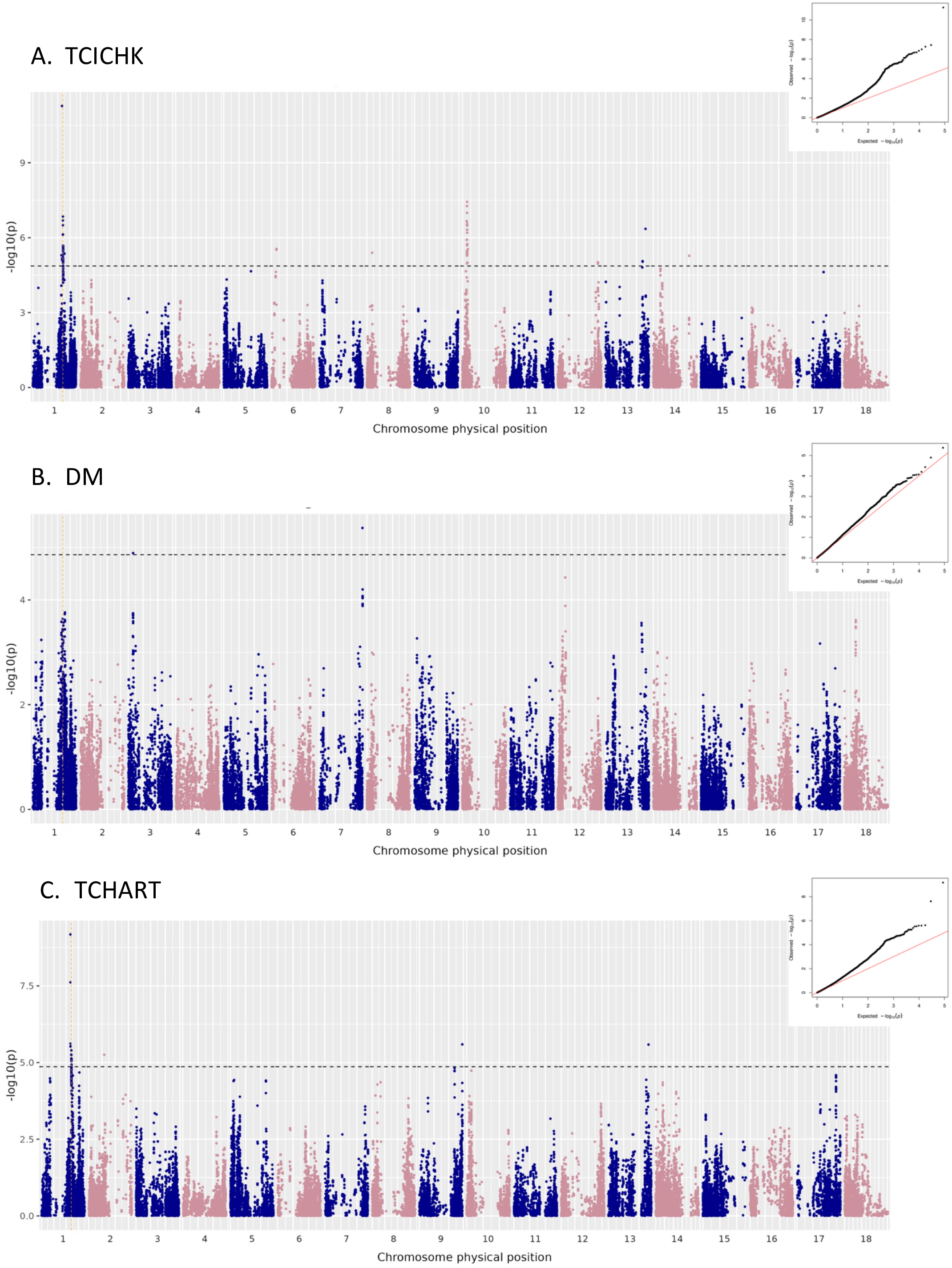
Manhattan plots for univariate GWAS for TCICHK (A), DM (B), and TCHART (C). The dashed black line indicates the modified Bonferroni significance threshold at α = 0.05. The vertical dashed orange line indicates the position of the *PSY2* causal locus used as a covariate. Q-Q plots for each analysis are at right.

### Candidate gene analysis

Annotated genes within a window of 100kb from the bivariate peak were identified as candidates based on the rate of LD decay. On most chromosomes, LD decayed to an average R^2^ of 0.25 at approximately 100kb physical distance, although chromosomes 1 and 4 had more extensive LD, likely due to the presence of large *M. glaziovii* introgressions on these chromosomes (Bredeson et al. 2016; Wolfe et al. 2019; Chan et al. 2022) (Fig. S6).

Candidate genes within 100kb of bivariate peak SNPs are listed in Table S5. None of the *a priori* candidate genes for the carotenoid or starch pathways were identified in this list. The peak SNP on chromosome 1 fell within the gene Manes.01G115400, annotated as a MYB-like DNA binding protein. The peak SNP on chromosome 10 fell within the coding region of Manes.10G034900, *ATP-binding cassette transporter B family member 20-related.* Candidate genes for other agronomic traits (DYLD, FRYLD, RTWT, HI, SPRTPERC, CMD3I, and CMD3S) from univariate analyses are listed in Table S6.

## Discussion

The main objective of this study was to test the hypothesis of mediated pleiotropy between TC and DM in cassava by examining genome-wide contributions to the negative covariance between traits. Bivariate mixed linear models were used to estimate the additive effects of each SNP on TC and DM while controlling for the effects of *PSY2* and genome-wide population structure. Four main points of evidence for mediated pleiotropy were considered: whether there are multiple QTL jointly associated with both traits, the extent of residual genetic covariance between traits after accounting for identified QTL, whether QTL tend to affect the traits in opposite directions, and whether QTL affect the traits in a direct or indirect manner.

The bivariate genome-wide associations identified two QTL passing the modified Bonferroni significance threshold at α = 0.05 on chromosomes 1 (S1_23484417) and 10 (S10_2963170) for joint association with TCICHK and DM. To our knowledge, the QTL on chromosome 10 jointly associated with TC and DM has not previously been reported. Rabbi et al. (2017) did show a single significant SNP associated only with TCHART on chromosome 10, but it does not appear to be in the same location as the QTL peak identified here. These associations therefore represent new observations of apparently pleiotropic QTL.

The incorporation of more diverse, early-stage selections within the yellow cassava breeding population likely facilitated the identification of these new loci. None of the TCICHK or TCHART univariate association peak SNPs overlapped with the DM univariate association peaks on chromosomes 3 and 7, despite the presence of the two significant bivariate TCICHK and DM peaks. This observation suggests that the increased power of the bivariate association model facilitated the identification of these joint associations which would have been missed if the two univariate associations were simply compared.

The estimated effect sizes of the chromosome 1 and 10 loci correspond to +2.28 *μ*g/g FW TC / -1.42% DM and -1.67 *μ*g/g FW TC / 0.92% DM, respectively (Fig. S3). While these effect sizes are large enough to potentially warrant marker-assisted selection strategies for individual traits, since these loci affect the traits in opposite directions, a genomics-assisted breeding method incorporating information from many markers would likely be more useful. In addition, the frequencies of the favorable TC allele at both loci are already quite high, consistent with the history of selection for yellow color in this breeding population (Fig. S7).

An additional 15 loci passed the relaxed significance threshold of p < 0.001 for joint association with TCICHK and DM. After accounting for all 17 of the peak SNPs at these loci and *PSY2* as covariates in a linear model, there was still statistically significant negative genetic covariance remaining between TCICHK and DM. With the putative QTL and *PSY2* accounting for 14.2% of the genetic covariance in total, the polygenic background still explained the majority of the covariance. Therefore, we conclude that there are significant polygenic contributions to the genetic correlation between TCICHK and DM, disrupting the hypothesis that *PSY2* is the main driver of the relationship.

We then tested whether QTL are more likely to contribute negatively to the covariance between traits than would be expected by chance. Under the hypothesis of a trade-off for carbon precursors between starch and carotenoid synthesis, any variants upregulating metabolic flux towards one pathway would be expected take away from the other pathway. Though the 17 peak SNPs used in the binomial test are not exactly independent (all SNPs were identified in only one dataset), we treated them as such given that they were mostly on different chromosomes and had low R^2^ with each other (R^2^ ≤ 0.095). In setting a more relaxed threshold for analyses on effect direction, we assumed that any false-positive peaks would be equally likely to affect the traits in the same or in opposite directions and hypothesized that an inherent biologically-driven trade-off would result in mostly opposite-direction effects.

While most peak SNPs (11 of 17) affected the traits in opposite directions, a binomial test of effect directions indicated that this pattern was not statistically significant. Therefore, while the overall genetic covariance is strongly negative, we cannot conclude an inherent underlying trade-off drives consistent opposite-direction effects across all QTL. The identification of several loci that affect the traits positively in the same direction, and variation among the magnitude of opposite-direction effect sizes such that not all effects were equal and opposite, suggests potential for making genetic gain on both traits. However, since none of the peaks affecting traits in the same direction passed the high-confidence significance threshold, we cannot recommend them as targets for marker-assisted selection unless further research can validate their effects. Rather, genomic selection leveraging many small effects throughout the genome may be a more effective approach.

To infer whether these joint associations were direct or mediated, we used model comparisons to test whether jointly-associated SNPs retained their associations with each individual trait after accounting for the other trait as a covariate. Both the chromosome 1 and 10 QTL peak SNPs retained significance in the univariate model for TCICHK after accounting for DM, but not in the univariate model for DM accounting for TCICHK. This result suggests that the QTL are more likely to be directly associated with TCICHK and indirectly associated with DM. Among all the suggestively-associated SNPs, the model comparison analysis supported a direct relationship with TCICHK for 9 SNPs and a direct relationship with DM for 4 SNPs, while no SNPs seemed to be directly associated with both traits simultaneously. These results support the hypothesis of mediated rather than direct pleiotropy (Fig. 1). Though the directionality of the mediating mechanism remains unclear, there is stronger evidence for TCICHK being the mediating trait since the majority of peak SNPs support that model. In addition, these data would support the hypothesis of a pleiotropy mechanism mediated by a latent variable such as a phytohormone signal, in which case it would be difficult to distinguish between direct/indirect and indirect/direct statistical associations (Fig. 1) (Stephens 2013).

Similarly, the lack of *a priori* candidate genes identified among the significant bivariate GWAS hits supports the hypothesis that the interaction between carotenoids and starch is mediated indirectly through regulatory pathways rather than direct pleiotropic action of genes in the biosynthetic pathways or direct competition for precursors (Villwock et al. 2024). The two previously identified candidate genes for DM on chromosome 1, *sucrose synthase* and *ADP-glc PPase*, did not show up as GWAS hits in this study, either because they were not causally polymorphic or because their effects were statistically eliminated by the *PSY2* covariate used in the association models.

Of the identified candidate genes (Table S5), several have potential biological relevance to the TCICHK and DM relationship. The gene containing the chromosome 1 peak SNP at S1_23484417, Manes.01G115400, is annotated as homologous to *Arabidopsis MYB4* (Table S5). In *Arabidopsis, MYB4* has been shown to affect leaf carotenoid content via transcriptional regulation of photooxidative stress response (Agarwal et al. 2020; Banerjee et al. 2024). While it is unclear whether Manes.01G115400 has an analogous function to *Arabidopsis MYB4,* other MYB-family transcription factors play roles in regulating carotenoid content as well (Sagawa et al. 2016; Zhu et al. 2017; Song et al. 2023).

At the chromosome 10 peak, the *ABC transporter* candidate gene is relevant to the hypothesis that ATP import into the amyloplast is a limiting factor for both starch and carotenoid synthesis (Villwock et al. 2024). Upstream 9.7kb from the chromosome 10 peak is *PsbP-like protein 2*, a chloroplastic protein involved in alternative electron transport to protect against photooxidative stress (Ishihara et al. 2007). In addition, an *AQUAPORIN/tonoplast intrinsic protein* (Manes.10G035000; *MeTIP1;4*) 21kb from the chromosome 10 peak SNP, is involved in vacuolar water and small solute transport (Zou and Yang 2019). Water content is an indirect component of both TC and DM measured per unit fresh weight. In *Arabidopsis, TIP3* expression is induced by an ABA-responsive transcription factor (Mao and Sun 2015). Therefore, *MeTIP1;4* could hypothetically be involved in increasing water uptake in response to higher ABA synthesis downstream of the carotenoid pathway, resulting in lower dry matter percentage. Candidate gene validation is needed to test these hypotheses in cassava and other starchy crops like sweetpotato exhibiting similar negative genetic relationships between TC and DM (Villwock et al. 2024). Future research will examine the co-expression patterns of candidate genes in cassava roots.

While the chromosome 1 QTL at 1_23484417 is relatively near the *PSY2* causal variant at 1_24155522, it appears to be a distinct QTL. Since the *PSY2* causal SNP (1_24155522) was imputed with a DR2 value of 0.74, it is possible there were imputation errors such that this covariate did not fully account for the *PSY2* effect. The *PSY2* SNP also explained a smaller proportion of the genetic variation in TCICHK (2.4%) than expected based on previous studies. Other methods of accounting for the *PSY2* effect in the association models were tested, including by fitting the closest non-imputed SNP to 1_24155522 as a fixed effect covariate, using the *PSY2* region (within 20kb of the coding region) haplotype as a random effect factor covariate, and using the first three PCs from a PCA of SNPs in the *PSY2* region as fixed effects. None of these methods significantly affected the results of an additional peak on chromosome 1 (data not shown). In addition, the 1_23484417 SNP was in low LD with SNPs in the *PSY2* region (Fig. 5). Therefore, we assume that the effect of the 1_24155522 variant in *PSY2* was effectively captured in the model. Since the new chromosome 1 QTL is 690kb upstream of the *PSY2* gene, it is possible that this QTL is capturing a *cis-*regulatory element of *PSY2.*

While imputing low-density genotype data to higher SNP density increases genetic mapping resolution, it also introduces imputation error. Using imputed genotype data has become standard practice in many genomic prediction and selection pipelines, in which low-cost genotyping assays are needed to screen large breeding populations, and is becoming more common in GWAS as well (Niehoff et al. 2022; Zhao et al. 2025). Using imputed genotype data has been shown to increase statistical power despite the addition of noise due to imputation error (Guan and Stephens 2008). To mitigate loss of statistical power due to imputation error, we retained the allele dosage estimates as decimals rather than rounded integers to help account for the uncertainty due to imputation error (Kutalik et al. 2011; Auer et al. 2023). Nevertheless, individual imputed marker-trait associations should be interpreted with caution and genotyped directly for QTL validation in future studies.

Q-Q plots from bivariate genome-wide associations with TCICHK and DM showed acceptable control of inflation from the genomic relationship matrix and the *PSY2* covariate. In the univariate analyses, there was considerable Q-Q plot inflation for TCICHK and TCHART. Although population structure and the *PSY2* were accounted for in these models, strong selection pressure for root yellowness as the primary trait in the yellow cassava breeding population likely created additional structure around causal loci. In practice, white-fleshed and yellow-fleshed cassava roots are easily distinguished, but quantitative variation among cream-colored roots blurs the line between them. After subsetting the data to retain only the yellow subpopulation, the positions of Manhattan plot peaks were similar, although with generally higher p-values reflecting lower power (data not shown). Therefore, it is unlikely that the major univariate QTL peaks for TCICHK and TCHART were driven by structure between the white and yellow subpopulations.

The precision of TCICHK phenotypes did not appear to outweigh the advantage of having many more TCHART observations, since TCHART had a higher H^2^ (0.73) compared to that of TCICHK (0.49). (Table 1). However, negative genetic relationships with yield traits (DRYLD, FRYLD, RTWT) were evident with TCICHK but not with TCHART. While TCICHK and colorimetric measurements are both highly correlated with HPLC measurements of TC (Jaramillo et al. 2018), TCHART may be influenced by other factors that affect visual perception of root color, while TCICHK may be more specific to carotenoids. The negative correlation between TCICHK and SPRTPERC (sprouting percentage; r = -0.25, p < 0.05) in these data is notable. This relationship has been previously observed in the field, though not formally described to our knowledge (P. Kulakow, personal communication, July 2021). We hypothesize that this negative correlation could be driven by the influence of carotenoid synthesis on downstream ABA, which regulates bud dormancy (Chen et al. 2020).

In summary, the presence of two new jointly-associated QTL identified at a high confidence level, residual negative genetic covariance after accounting for 17 putatively associated SNPs tending to affect the traits in opposite directions, and a consistent pattern of direct/indirect statistical associations with the two traits supports the hypothesis of polygenic, mediated pleiotropy driving the relationship between TC and DM. While some plausible candidate genes were identified, none were directly in the carotenoid or starch biosynthesis pathways.

Furthermore, there was not clear evidence for the direction of the mediation of one trait on the other, and one QTL affecting the two traits in the same direction was also identified. This suggests that the relationship between TC and DM is likely not the result of a simple trade-off or a single causal chain, but more complex regulatory interactions. In addition, new QTL for TC were identified that expand our understanding of the genetic basis of TC accumulation in cassava beyond the *PSY2* locus.

This study represents a new application for multivariate GWAS that goes beyond identifying individual variants to test hypotheses about traits’ genetic relationships across the genome. Our approach builds on previous multivariate GWAS methods that explicitly estimate SNP effects on each trait, which has a built-in interpretation framework, as opposed to the difficult-to-interpret single p-values given from an overall multivariate association. A downside to our method is that the computational time is high, and it may not be feasible to extend to more than two traits. More efficient but less flexible implementations similar to our multivariate GWAS method are available in the MTMM and GEMMA software (Korte et al. 2012; Zhou and Stephens 2014). Multivariate GEMMA output very similar GWAS results with the data used here and ran much faster, but GEMMA does not offer the same flexibility in specifying and analyzing covariance structures as ASReml-based models do. Future studies could improve on these methods by improving the computational time while retaining the modeling flexibility.

Overall, the identification of polygenic mediated pleiotropy in this study demonstrates a multivariate GWAS framework for investigating the genetic architecture of correlated traits, providing insights to guide both applied breeding strategies and future research into underlying biological mechanisms.

## Data availability

All datasets and R code used for these analyses are available on GitHub (http://www.github.com/serenvillwock/mvGWAS). Raw field trial data and imputed genotype files are additionally available on http://www.cassavabase.org/ under the public list ‘yellowGWAS23_trials_plusCASS’. The reference panel data used for genotype imputation is available at https://www.cassavabase.org/ftp/marnin_datasets/nextgenImputation2019/ImputationStageIII_72619/. *Manihot esculenta* v6.1 reference genome and gene annotations are available on Phytozome at https://phytozome-next.jgi.doe.gov/info/Mesculenta_v6_1.

## Acknowledgements

We are grateful to the Cassava Breeding Unit team at the International Institute of Tropical Agriculture, Ibadan, Nigeria, especially including Lynda Adaobi Nnadi, Adenike Damilola Ige, Peter Iluebby, Patrick Akpotuzor, Victor Monday Chukwuyem, Toye Ayankanmi, and Kehinde Nafiu for their assistance with tissue sampling and field trials; Prasad Peteti, Ekanem Ukoabasi, and Mahmud Kehinde Dhikrullah for their assistance with data curation; and Peter Kulakow, Peace Ikeanyi, and Richard Ofei for their administrative support. The development of the breeding populations used in this study was sponsored by HarvestPlus and the NextGen Cassava project. We are grateful to Kelly Robbins and to the Cornell University BRC Bioinformatics Core Facility (RRID:SCR_021757) for providing access to computational resources, Michael Gore for insightful discussions, and Salvador Gezan at VSN International for providing ASReml-R technical support. We thank Andrew Siefert for statistical consultation. The authors thank the UK’s Foreign, Commonwealth & Development Office (FCDO) and the Bill & Melinda Gates Foundation (Grant INV 007637; www.gatesfoundation.org) for their financial support. SSV is supported by the USDA National Institute of Food and Agriculture AFRI Predoctoral Fellowship project accession no. 1030847.

## Supplementary Information

**Figure S1.**
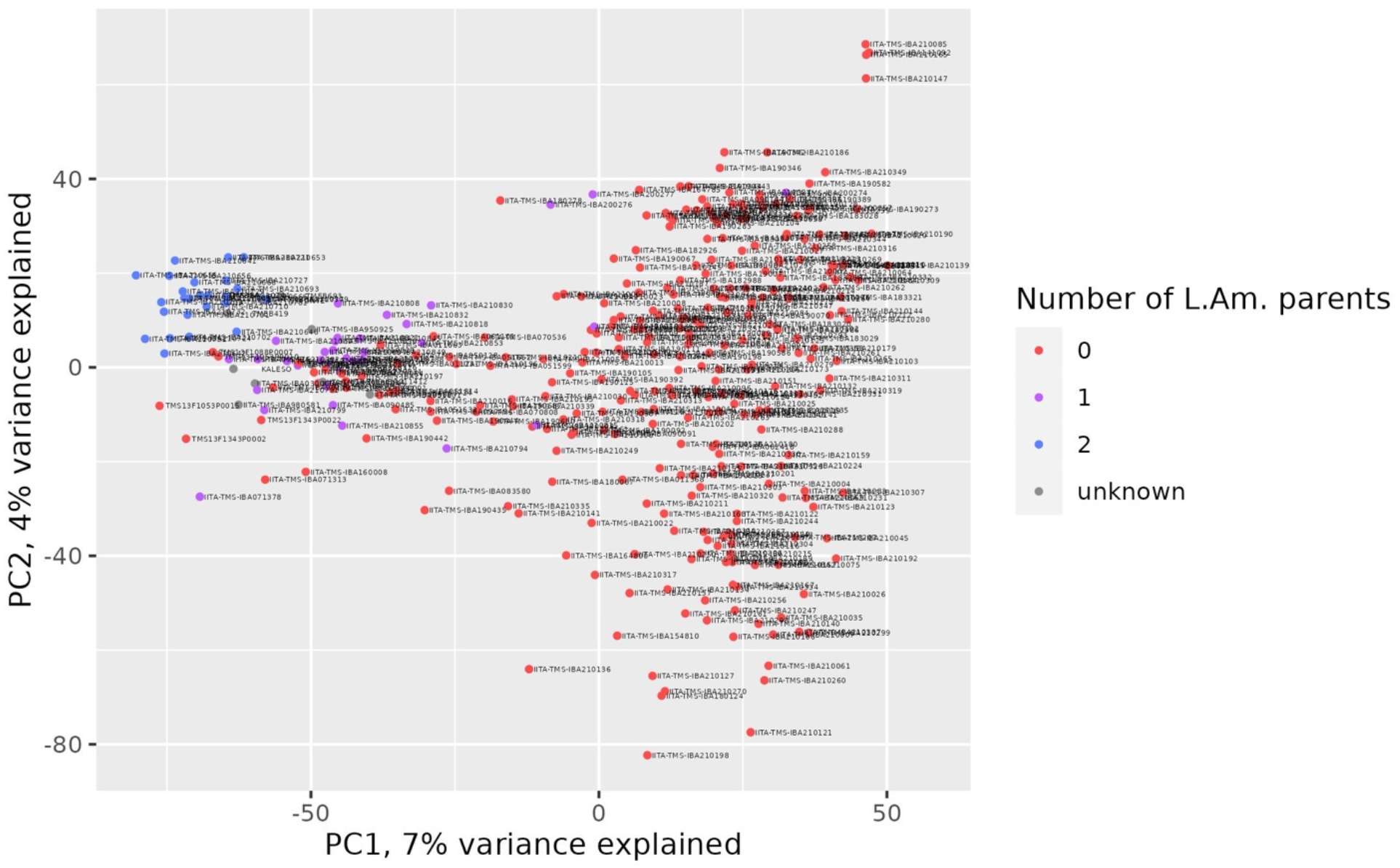
Principal component analysis of genotyped accessions shows that the first principal component (PC1) separates a cluster of accessions with recent Latin American ancestry. Color represents the number of Latin American parents listed in an accession’s pedigree.

**Figure S2.**
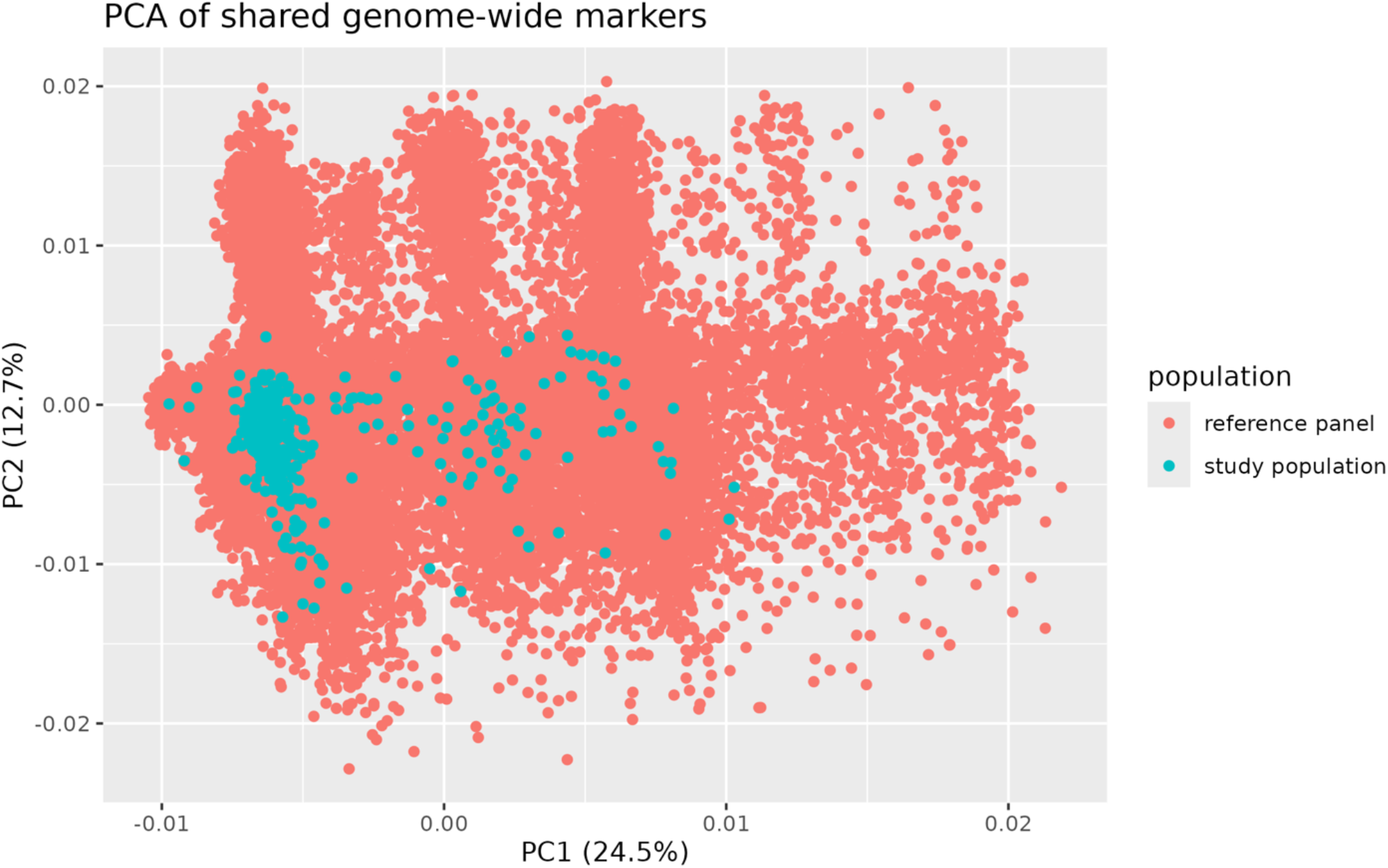
Principal component analysis of 2,939 genome-wide genetic markers shared across the imputation reference panel and the study population.

**Figure S3.**
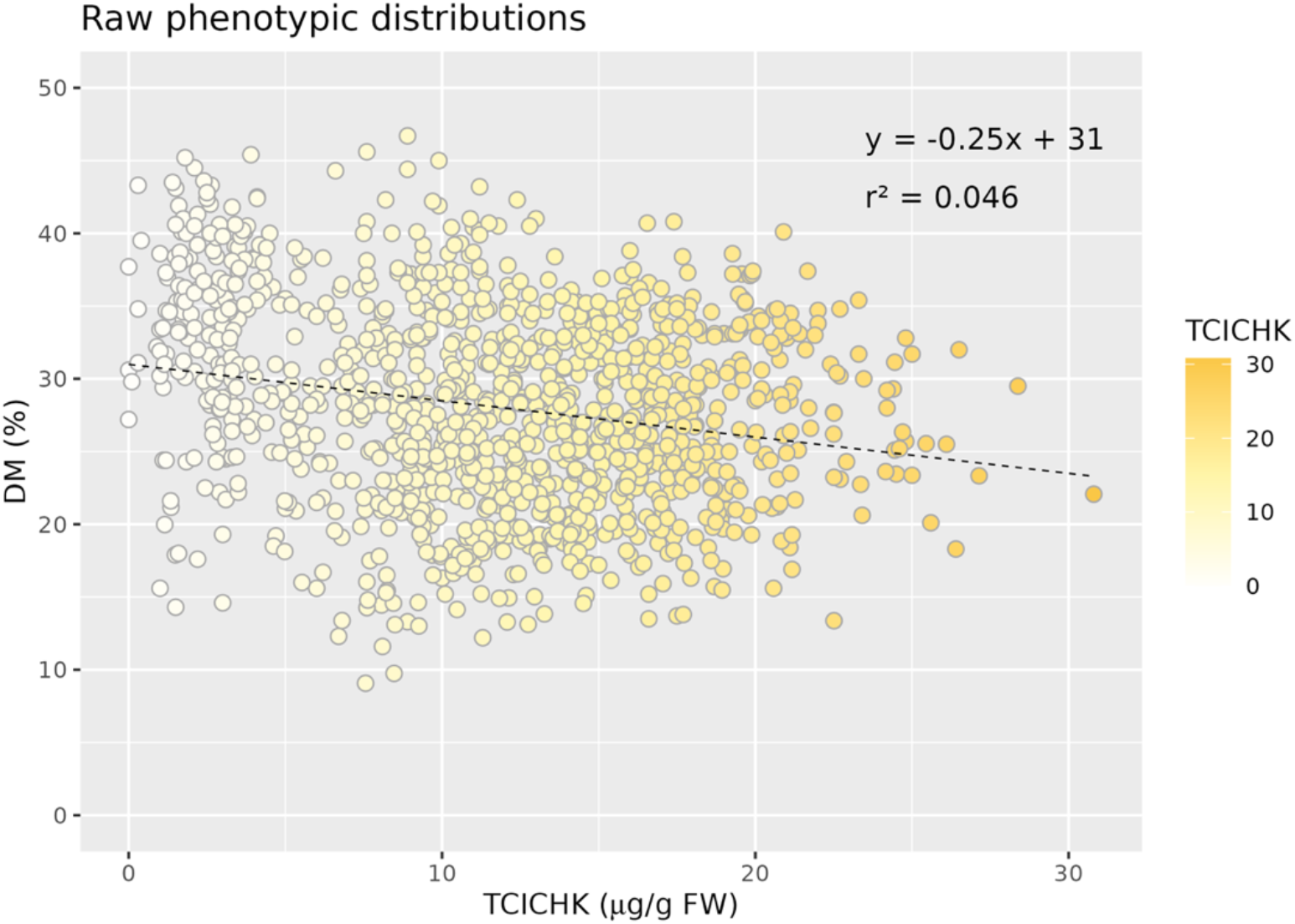
Raw phenotypic distributions of percent dry matter (DM) and total carotenoids measured by iCheck (TCICHK) in μg/g fresh weight. Points represent individual plot-level observations.

**Figure S4.**
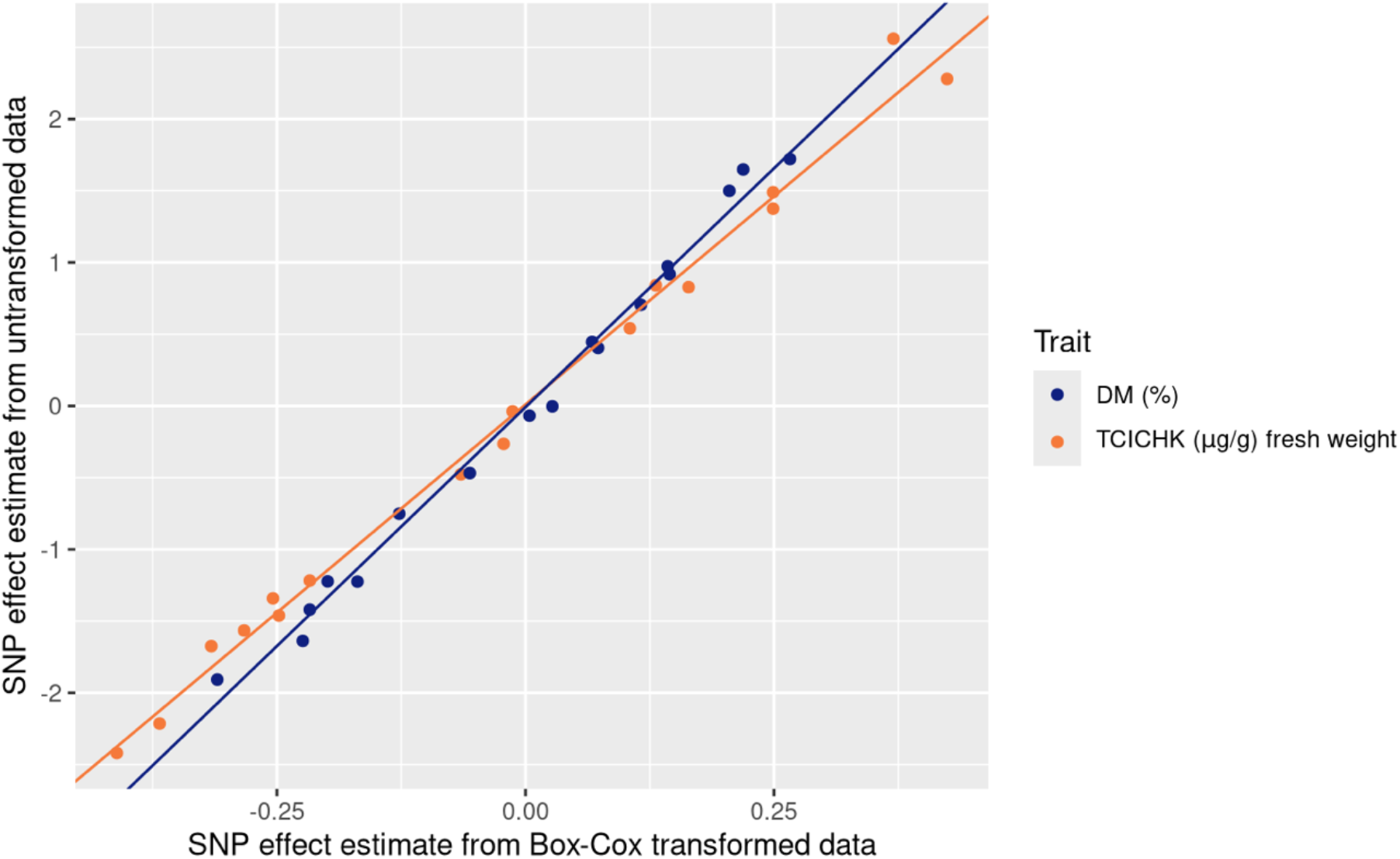
Comparison of peak SNP effect estimates from Box-Cox normalized data and from untransformed data show a linear relationship. SNP effect estimates from the untransformed data correspond to the traits’ original units.

**Figure S5.**
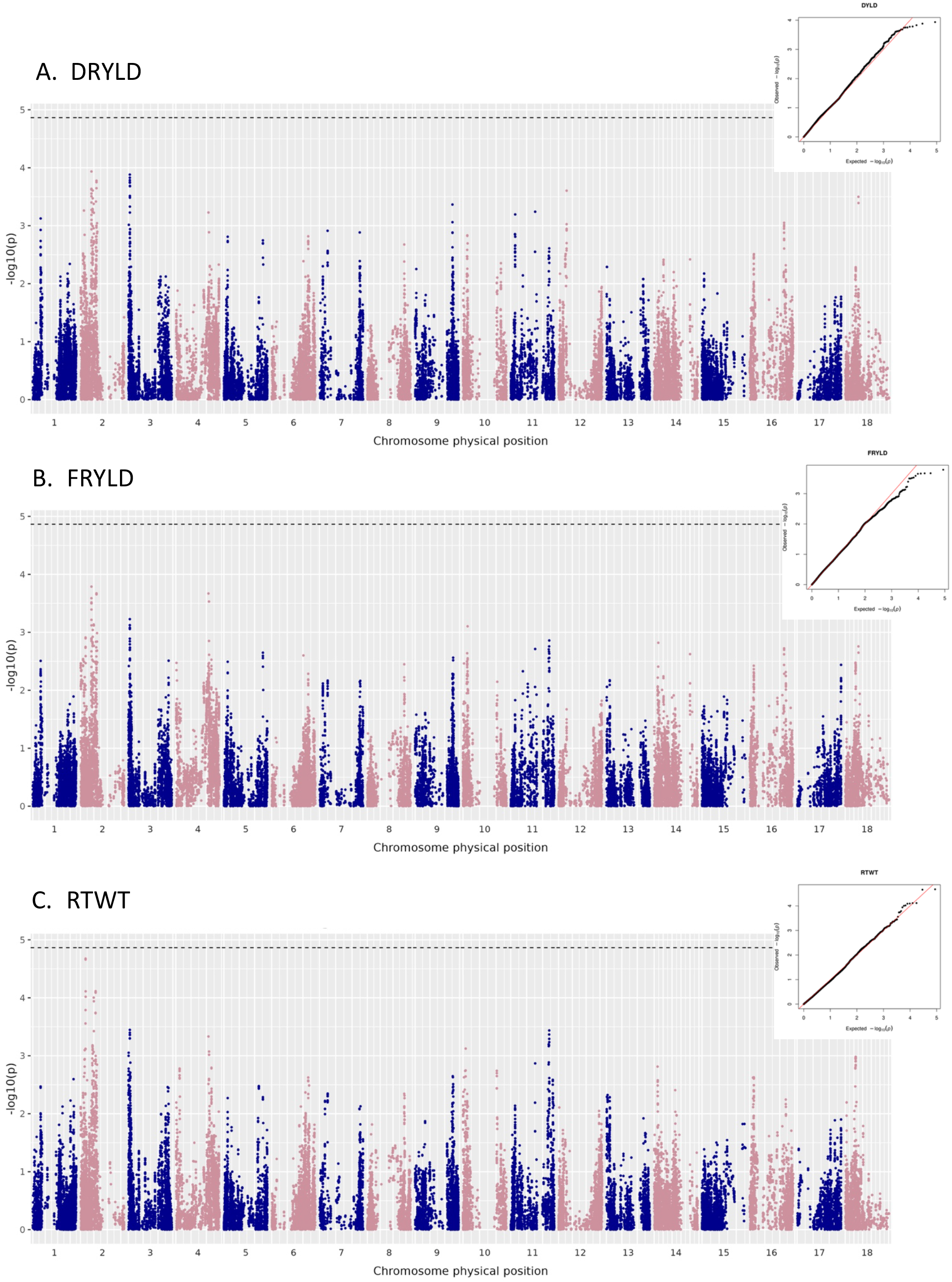
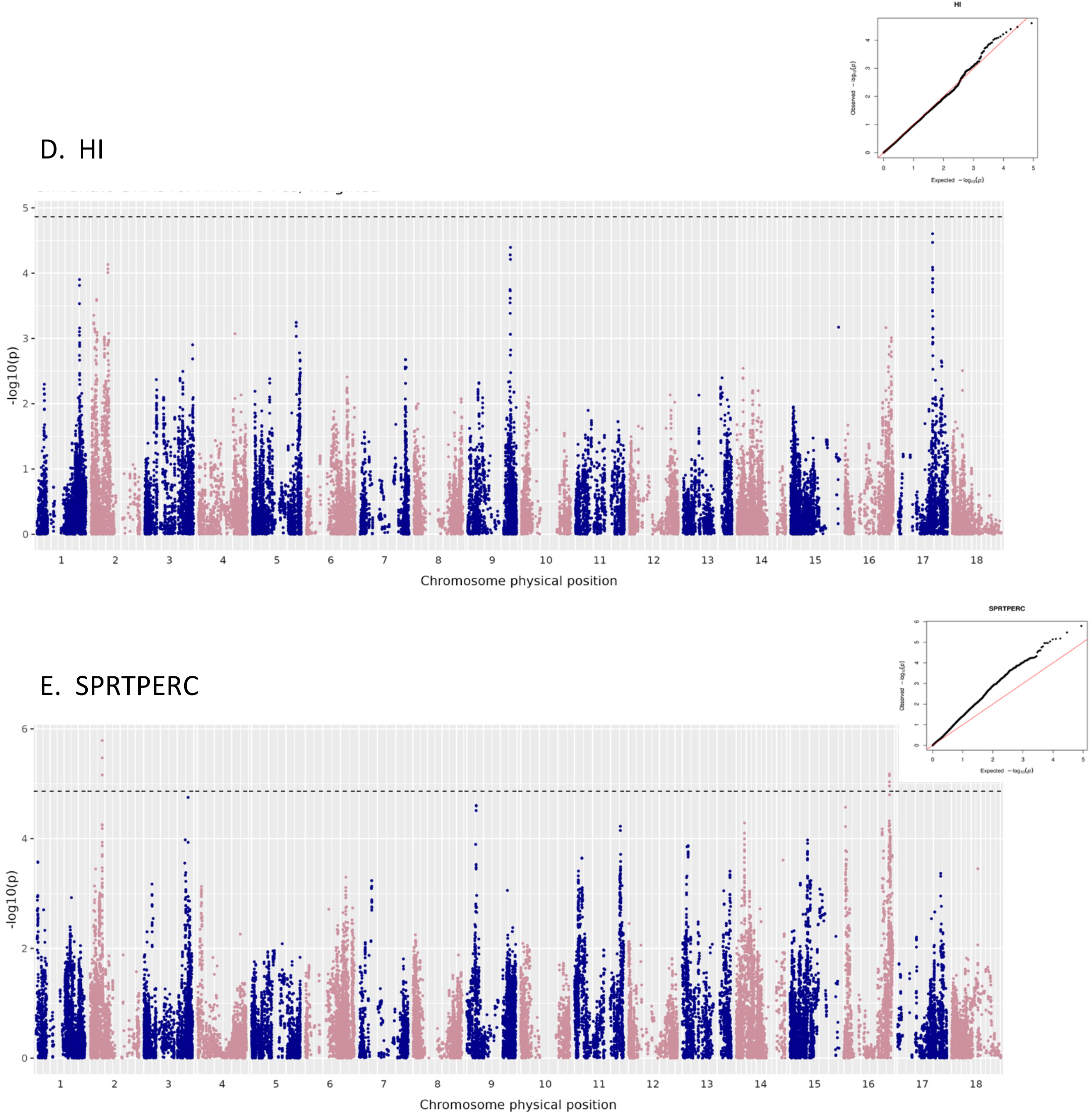
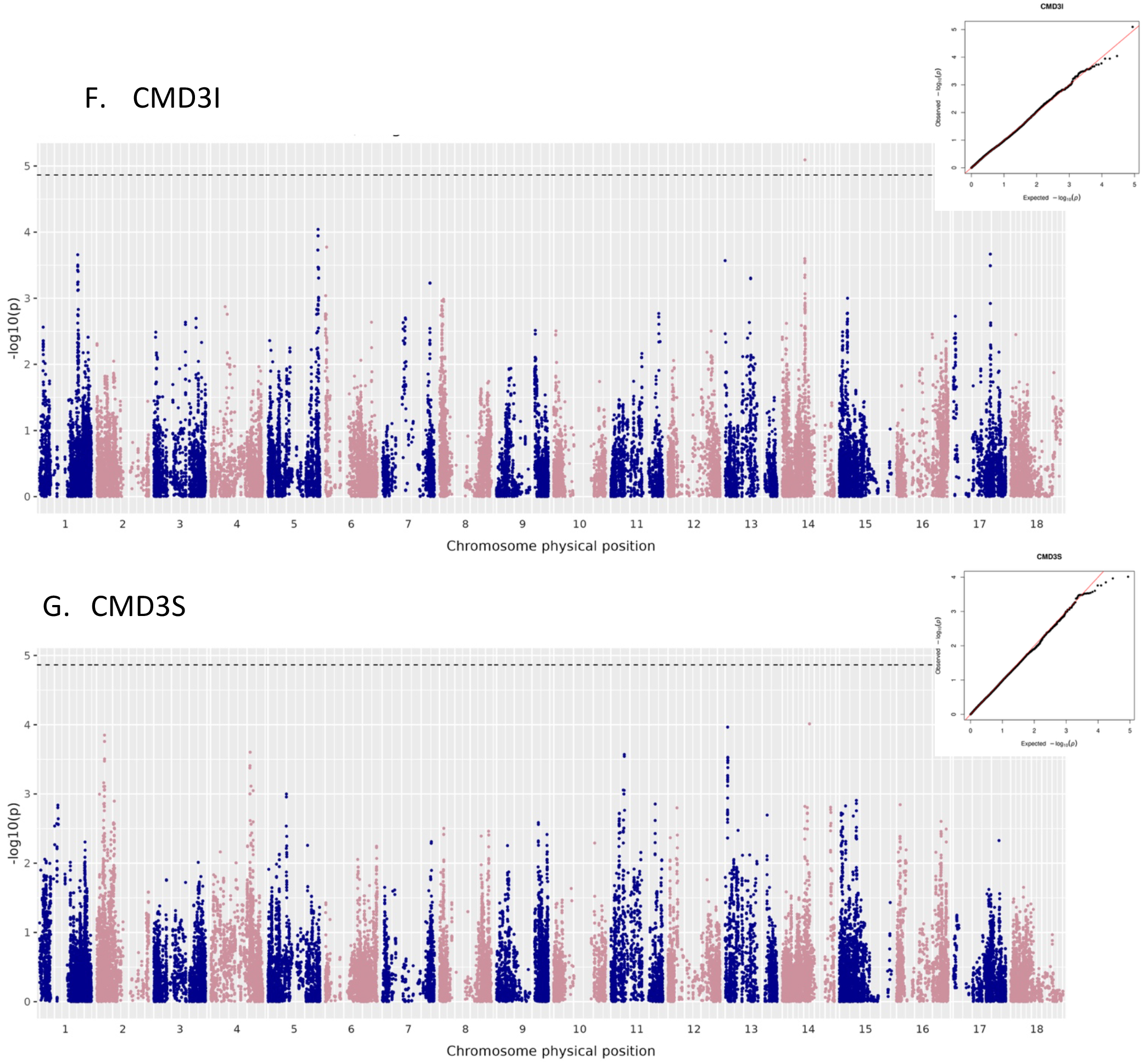
Manhattan plots for univariate GWAS for analyzed agronomic traits: DRYLD (A), FRYLD (B), RTWT (C), HI (D), SPRTPERC (E), CMD3I (F), and CMD3S (G). The dashed black line indicates the modified Bonferroni significance threshold at α = 0.05. Q-Q plots for each analysis are in the upper right corner.

**Figure S6.**
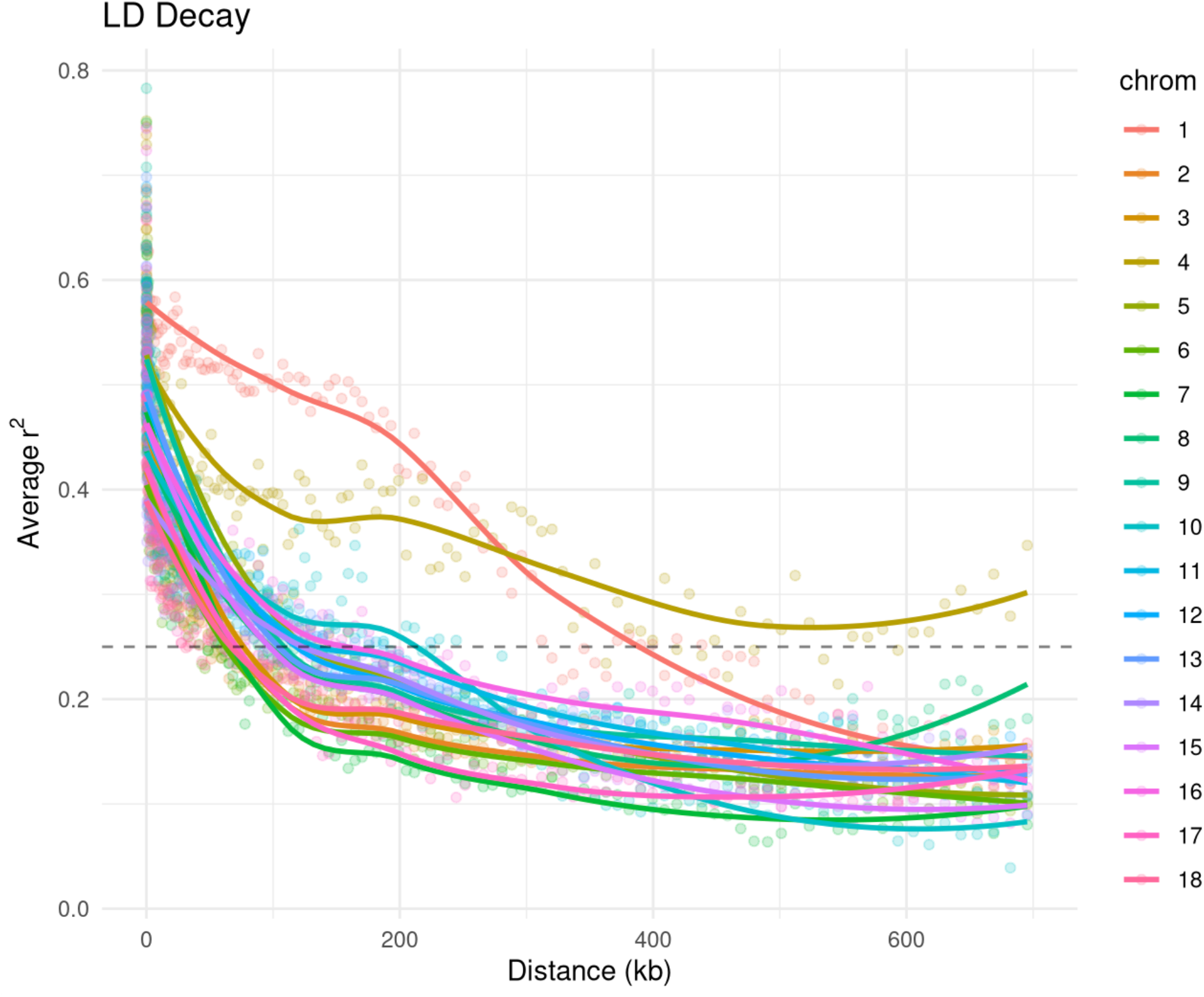
Linkage disequilibrium (LD) decay measured pairwise r^2^ between SNPs. Points represent average pairwise *r*^2^ per chromosome within 200 physical distance intervals.

**Figure S7.**
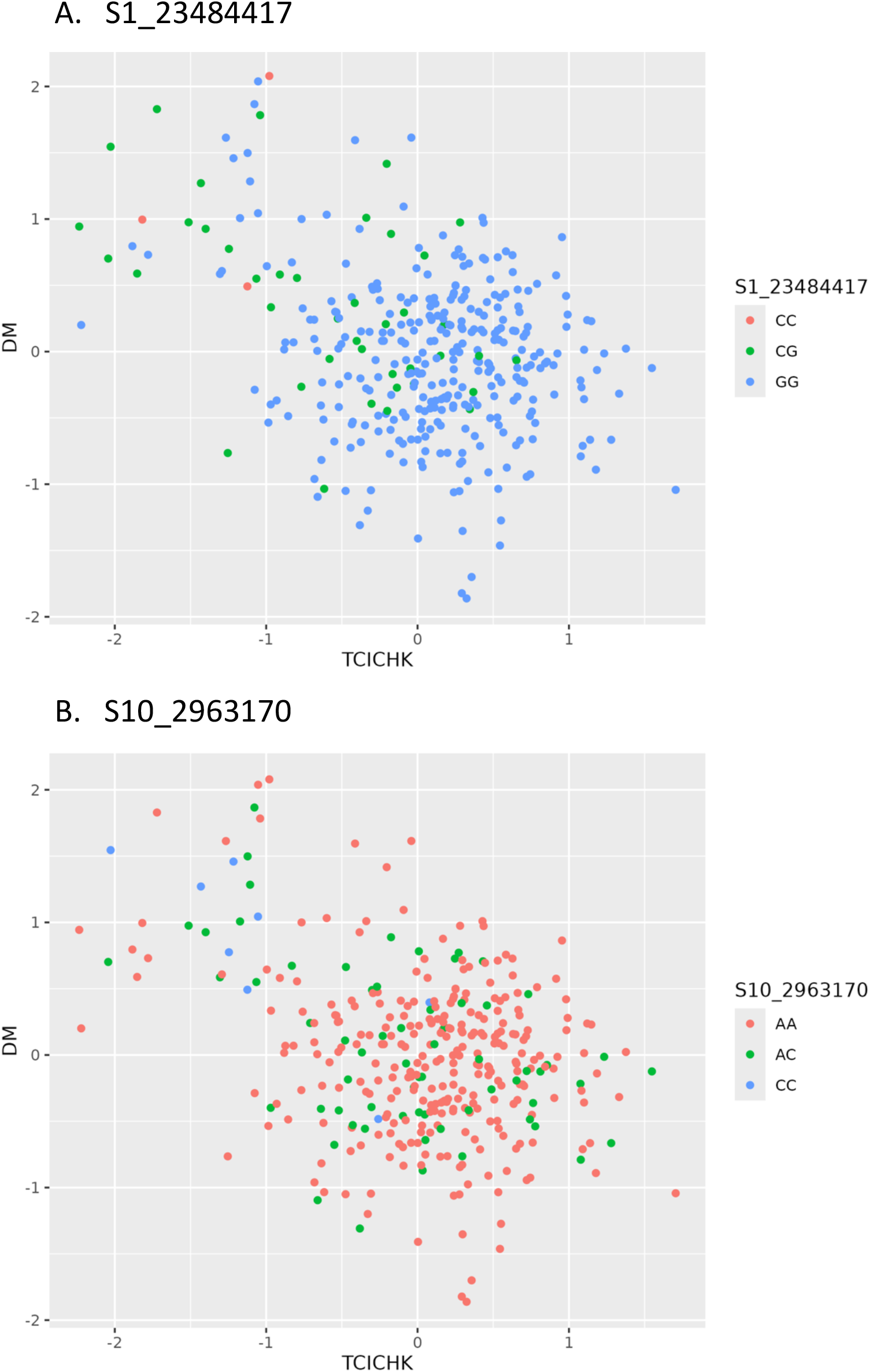
Genotype frequencies and phenotype distributions at the QTL peak SNPs identified on chromosomes 1 (A; S1_23484417) and 10 (B; S10_2963170). The x- and y-axes show the accession BLUP values for TCICHK and DM, respectively. Points are colored according to the alternate allele dosage.

## Supplementary Tables

*Attached file.*

**Table S1.** Metadata for cassava accessions included in this analysis.

**Table S2.** Metadata for field trials included in this analysis.

**Table S3**. *A priori* candidate genes with annotations relating to carotenoid, apocarotenoid, and starch synthesis and degradation pathways.

**Table S4.** Summary statistics for peak SNPs identified in univariate associations with agronomic traits at the Bonferroni-corrected significance threshold.

**Table S5.** Candidate genes within 100kb of peak SNPs significantly associated with TCICHK and DM at the modified Bonferroni significance threshold from the bivariate GWAS.

**Table S6.** Candidate genes within 100kb of peak SNPs identified in univariate associations with agronomic traits at the Bonferroni-corrected significance threshold.

